# Membrane proteins retain native architecture through native ESI and soft-landing

**DOI:** 10.64898/2026.01.13.699114

**Authors:** Jingjin Fan, Clare De’Ath, Lukas Eriksson, Carl von Hallerstein, Louise J. Persson, Abraham O. Oluwole, Noor Naseeb, Aziz Qureshi, Susanne Mesoy, Laurence T. Seeley, Simon B. Knoblauch, Neha Kalmankar, Erik G. Marklund, Tim Esser, Carol V. Robinson, Lindsay Baker, Stephan Rauschenbach

**Author notes:** Currently at Thermo Fisher Scientific, De Schakel 2, 5651GH Eindhoven, The Netherland (Tim Esser).

## Abstract

Native MS offers a clear picture of membrane protein stoichiometry and interactions, but it lacks direct structural insights at high resolution. Here, we examine the extent to which solution-phase structure and architecture can be retained after native, soft-landing electrospray ion beam deposition (ESIBD) by interrogating several membrane-protein complexes of diverse folds and oligomeric states by cryoEM. The overall protein architectures with secondary structure motifs can be observed after gas-phase transfer, soft landing, and embedding in amorphous ice. Notably, we determined the structure of the ammonium transporter AmtB at sub-3 Å resolution. It is nearly identical to the structure of the plunge-frozen control and even shows an extended C-terminal segment of AmtB, a dynamic region absent in the solution-phase structure. Our analysis shows that detergent adducts preserve membrane protein structure in vacuum by minimising destabilization of solvent-exposed regions and stabilization through additional polar contacts in vacuo. Molecular dynamics (MD) simulations support these results, suggesting that a monolayer shell of surfactant adducts avoids destabilization driven by unshielded polar residues and disruption of hydrogen bond networks. Overall, our findings provide a structural framework for integrating native MS with cryo-EM showing that gas-phase transfer and surfactant stabilisation preserves key architectural features and high-resolution structure of membrane proteins.

## Introduction

Membrane proteins (MP) comprise over half of all drug targets due to their central roles in cellular physiology, mediating critical processes such as selective transport of ions and molecules, signal transduction, and lipid metabolism.(*1, 2*). To study the functions of MPs in relation to their structure and environment, native MS has become an essential tool, capturing stoichiometries, lipid interactions, and drug binding states that often elude conventional structural methods.(*3–5*) In particular, when combined with complementary, structurally sensitive approaches such as ion mobility spectrometry(*6–8*), hydrogen-deuterium exchange (HDX)(*9, 10*) or molecular dynamics (MD)(*11, 12*), native MS offers unique capabilities to analyse heterogenous samples and capture molecular shapes, interactions and conformational variability. This can illuminate variable effects on binding sites, protein stability, and stoichiometry all while requiring minimal sample material. (*7, 13–15*) However, an enduring question remains unresolved: to what extent do MPs retain their native structures after transfer into the gas phase via native electrospray ionization? (*16–18*) This uncertainty, unresolved by direct structural evidence, complicates the interpretation of native MS data (*19, 20*).

High-resolution structure determination of MPs reconstituted in detergent micelles or other mimetics is routinely performed by X-ray crystallography or cryo-electron microscopy (cryoEM). (*21, 22*) Yet, selecting suitable mimetics(*22, 23*) and defining ligand/lipid binding stoichiometry(*24*) remain major challenges. Native MS can provide insight on both these issues, (*25*) but as the sample preparation is fundamentally different from crystallography or cryoEM, questions remain about the relevance across methods.

Soft-landing electrospray ion beam deposition (ESIBD) provides a route to molecular structure by gently depositing mass-selected large molecules onto sample carriers under vacuum.(*26–29*) After transfer into the gas phase and landing, soluble proteins and even viruses have previously been demonstrated to retain their overall shape when imaged by electron microscopy or holography (*30–32*) and in some cases show biological activity (*33–36*). Coupling ESIBD to high-resolution cryoEM imaging has enabled resolution of gas phase protein secondary structure and side chains (*37, 38*). These experiments support the growing view that native-like conformations can be preserved in vacuum.

Membrane proteins pose additional challenges. Their dependence on the membrane environment raises additional questions about conformational stability in the gas phase after the removal of membrane mimetics and dehydration.(*7, 39*) No method has yet resolved gas-phase prepared MP structures, leaving conformational preservation speculative despite hints that aspects of the native state may survive.(*19, 40*)

To address this challenge, we developed an ESIBD workflow for membrane proteins applied to multiple complexes. We mitigate unscreening of electrostatic interactions and exposure of hydrophobic patches due to dehydration, by retaining detergent, controlled via MS. Molecular dynamics (MD) simulations, when compared to the experimental results, reveals preserved overall architecture, protein-dependent behavious and small dehydration-induced changes. Under optimal detergent conditions, cryoEM imaging of soft-landed ammonium transporter B (AmtB) is achievable by embedding the protein in a vitreous layer of ice and produces a high-resolution density map with a resolution better than 3 Å.

## Results and Discussion

### Electrospray Ion Beam Deposition

Deposition and imaging of membrane proteins was performed by combining native ESIBD with cryoEM on a platform previously established for soluble proteins (*37, 38, 41*) (see **Methods** and **Figure S1** for details). We selected four membrane protein complexes spanning various oligomeric states (trimer, tetramer, pentamer), secondary structures (α-helices, β-barrels, β-sheets), and domain compositions (transmembrane and soluble): ammonium transporter B (AmtB), aquaporin Z (AqpZ), outer membrane protein F (OmpF) from *Escherichia coli*, and the *Dickeya dadantti* (previously *Erwinia chrysanthemi*) ligand-gated ion channel (ELIC). After gentle ionization by native ESI, the number of detergent molecules surrounding each protein was modulated by activation. Membrane protein ions were landed onto a freestanding carbon film held at cryogenic temperature, ensuring immobilization on the surface (*30*). A thin layer (∼20 nm) of amorphous ice was grown after landing the proteins. (*38*) The cryo-shuttle containing the vitrified grid was then transferred from the vacuum chamber into liquid nitrogen via a dry nitrogen, purged system and subsequently to the microscope under liquid nitrogen. The workflow is shown schematically in **Figure 1A**. Samples without an amorphous ice layer (called ‘ice-free’ henceforth) were also produced to represent adsorbed gas-phase proteins.

**Fig. 1.**
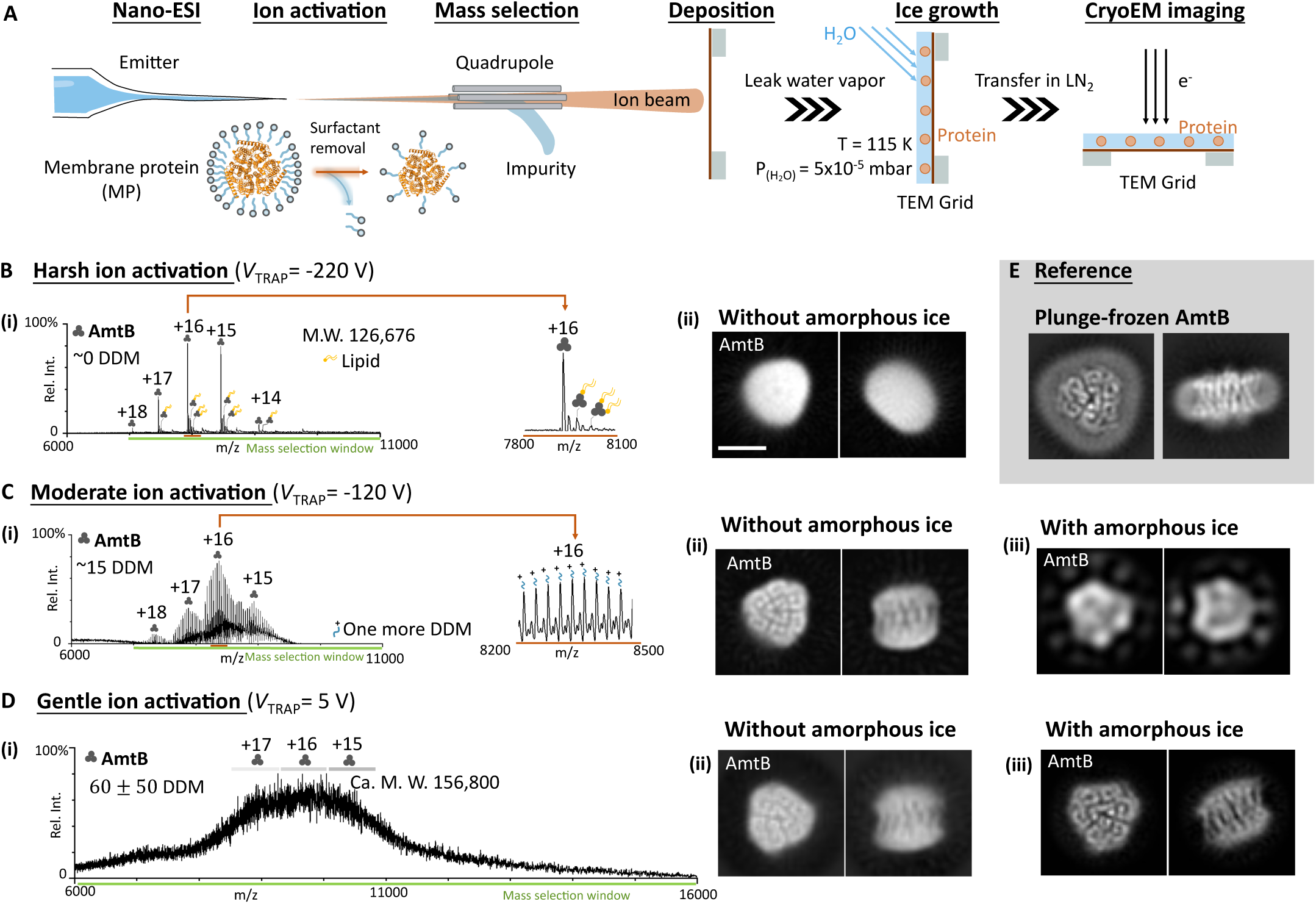
Activation-dependent detergent binding in the mass spectra of AmtB, and the corresponding cryoEM 2D classes acquired with and without growth of amorphous ice. (**A**) Schematic of ESIBD cryoEM sample preparation workflow, including electrospray ionization, detergent removal, mass selection, cryo-deposition, growth of amorphous ice, and cryoEM imaging (see **Methods**). **(B-D)** Mass spectra and corresponding cryoEM 2D classes of AmtB under varying ion activation conditions: harsh, moderate, and gentle. (**i**) Mass spectra and representative 2D classes (**ii**) without and (**iii**) with amorphous ice embedding (see **Figure S4** for detailed 2D class averages). (**E**) Reference 2D class averages of plunge-frozen AmtB. All 2D class averages here are shown at the same magnification; scale bar, 5 nm.

### Ion activation and detergent retention affects membrane protein structure

In native MS, collisional activation is often used to improve spectral resolution by removing adducts such as salts, water, or detergents. For membrane proteins, charge states typically correspond to varying numbers of detergent as well as lipid molecules bound to the protein.(*3, 42*) Different activation conditions, defined by the magnitude of a trapping potential (*V*_TRAP_), and catagorised as harsh, moderate, and gentle, are employed to alter the amount of bound detergent. (see **Figure S2** for details)

The impact of detergent binding and activation on membrane protein structure, is studied in detail for two oligomeric membrane proteins AmtB (M. W. = 126 kDa, **Figure 1**) and AqpZ (M. W. = 98 kDa, **Figure S3**). Under harsh activation conditions (*V*_TRAP_ = -220 V), the mass spectra exhibit sharp peaks, indicating effective removal of detergent though collisions. Additional low intensity peaks suggest the presence of a few bound lipids. Following mass selection, charge states from +18 to +14 of AmtB (**Figure 1B i**.) are deposited, and imaged. The resulting 2D class averages from cryoEM reveal the approximate shape of AmtB (**Figure 1B ii**.), but no structural detail.

When the trapping potential is decreased (moderate activation, *V*_TRAP_ = -120 V), the mass spectra display many adduct peaks for each charge state, corresponding to the retention of approximately 15 DDM detergent molecules, (**Figure 1C i**.). Corresponding ice-free 2D class averages show significantly improved resolution compared to the harsh conditions above, with nanometer-scale helical structural features visible in both top and side views (**Figure 1C ii**.). After embedding in amorphous ice, however, only the approximate shape of AmtB is observed, secondary structure is lost, likely due to increased structural heterogeneity induced upon the growth of the ice layer (**Figure 1C iii**.).

Under gentle activation (*V*_TRAP_ = +5 V), no additional trapping potential is applied, and protein ions are transmitted with a substantial number of associated detergent molecules. This condition results in a characteristic broad feature in the mass spectra **(Figure 1D i**.), consisting of many overlapping peaks of AmtB bound to different number of detergent molecules. Systematically varying the activation potential reveals that only AmtB with bound detergent persists in the ion beam, enabling quantification of the detergent that remains as a function of the applied activation (**Figure S2B**). We estimate that approximately 60 ± 50 DDM adducts are present per AmtB trimer under gentle activation conditions, which would comprise approximately one molecular monolayer on the protein surface (see **Figure S5).** 2D class averages from gentle activation with and without amorphous ice both show clear structural detail including well-resolved helical features (**Figure 1D ii., iii**.). Interestingly, no detergent micelle is visible, in contrast to the equivalent plunge-frozen AmtB, where the detergent micelle is a clearly visible feature (**Figure 1E**). Similar effects were observed for AqpZ using the same procedure, as shown in **Figure S3**.

These results show two key factors to obtain high-resolution cryoEM data from MPs: maintence of the membrane mimetic and embedding in amorphous ice. Increasing detergent retention by minimizing activation enhances structural preservation but a full micelle encapsulation is not necessary to stabilize AmtB and AqpZ during gas-phase transfer. Subsequent amorphous ice embedding together with a monolayer of detergent molecules is sufficient. At medium ion activation, the small number of detergent molecules (approx. 15 DDM) is sufficient to stabilize the MP during ionization and in the gas-phase. The loss of resolution upon water-ice embedding, implicates the protein-water interfaces as a driving force behind structural heterogeneity.

### Mechanism of detergent aided structural preservation

To rationalise the structural preservation of membrane proteins in the ESIBD process, we considered the role of detergent-protein interactions in vacuum using our experimental data and MD simulations. Extensive work has already characterised how ion–gas collisions and ion activation affect protein structure in MS(*43–45*), so here we focus specifically on the consequences of water and detergent removal. Starting from an equilibrated solution-phase system of membrane proteins with 150 detergent molecules, we initiated vacuum simulation with 0, 15, and 150 detergent molecules per protein, mirroring the conditions after harsh, moderate and gentle activation in native MS.

MD models for 0, 15, and 150 detergent molecules per AmtB trimer, coloured by Root-mean-square-deviation (RMSD) relative to the solution-equilibrated structure, reveal that the deviations occur at the rim of the protein molecule (**Figure 2A**). A calculation of the local solvent-exposure (SE) reveals the same pattern. For each residue, SE evaluates the amount and proximity of solvent interaction versus protein or detergent interactions within a 5-nm range. High exposure scores correspond to highly solvent-exposed regions (**Figure S6,** further details in SI **Methods** and the previous paper(*38*)), which reveals the sites that are most affected by dehydration (*46, 47*). Solvent exposure mapped for AmtB bound to **(i)** 0, **(ii)** 15, and **(iii)** 150 DDM closely tracks by-residue backbone RMSD-coloured models. **This local picture is mirrored at the global level;** backbone RMSD of AmtB across trajectories clearly shows the stabilising effect of detergent, increasing from 3.2 Å for 150 DDM to 4–5 Å and 5–6 Å for systems with 15 or 0 DDM, respectively (**Figure 2B**, and **Table S1**). Simulation and SE results are thus in accordance with previous experimental observations showing a loss of secondary structure resolution as detergent is removed and further reveal that peripheral regions are most susceptible to vacuum induced distortions in the absence of detergent.

**Fig. 2.**
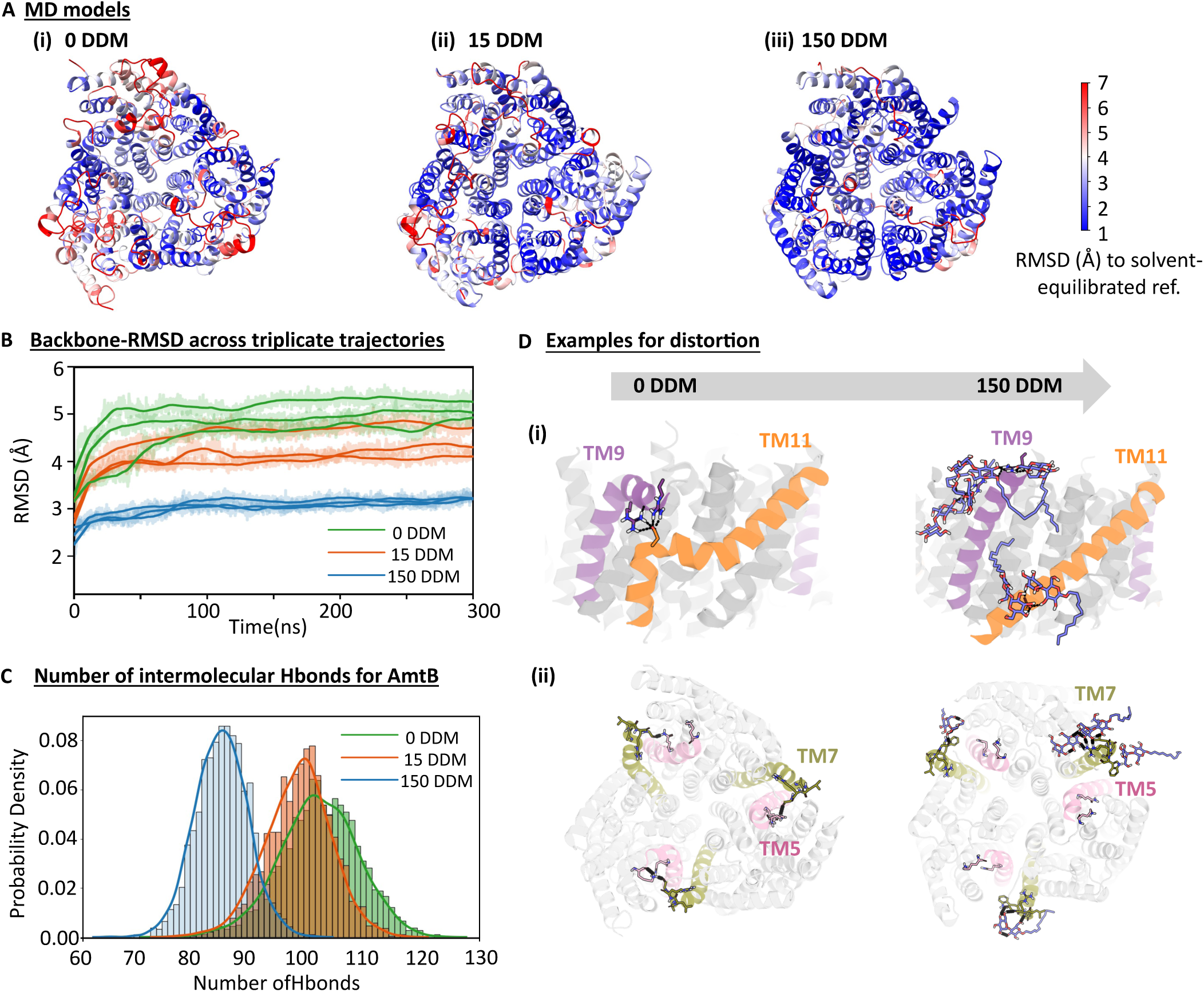
Molecular dynamics simulations of AmtB with DDM molecules in vacuum. **(A)** Models representative for each condition coloured by per-residue back-bone RMSD (blue, low RMSD; red, high RMSD). Detergent molecules are hidden. **(B)** Backbone-RMSD across triplicate trajectories show minimal deviations with 150 DDM (∼3 Å) but larger deviations with 15 DDM (4-5 Å) or no DDM (5-6 Å). **(C)** Distribution of intramolecular hydrogen bonds of AmtB across detergent levels. **(D)** In the absence of detergent, helices in AmtB are distorted, for example, (i) increased bending of peripheral helices TM9 and TM11 and (ii) surficial residues of TM5 and TM7 also form non-native contacts. Addition of detergent mitigates these distortions.

MD simulations and SE consideration also allowed us to shine a light on the mechanism by which detergent achieves this mitigation of dehydration effects. SE calculations suggests that detergent adducts provide contact sites for peripheral residues which would otherwise be directly exposed. Though van-der-Waals contacts between the hydrophobic detergent tails and the TM-region may contribute to stabilisation, MD simulations suggest that the polar headgroups of DDM (two saccharide rings) take on a particularly important role by serving as binding partners to polar sidechains and thus mitigating dehydration-induced distortions. (*48, 49*) In solution, water forms hydration shells around polar and charged residues, while detergent covers hydrophobic surfaces, together supporting folding and stability. However, in vacuo, the absence of solvent detergent leaves electrostatic interactions unscreened and exposes hydrophobic patches. (*16*) MD simulations show that in the absence of detergent, polar sidechains of the MP replace sidechain-solvent interactions with sidechain-sidechain hydrogen bonds, a phenomenon likely enhanced by the absence of screening of Coulombic interactions in vacuum. (*50*) The protection of detergent is consistent with previous simulation-based predictions of detergent function in vacuum.(*11, 12, 49, 51*) Quantifying the number of intra-molecular (i.e. within complex) hydrogen bonds via MD between sidechains of AmtB for different amounts of detergent show this behaviour (**Figure 2C)**. Helical distortion is one possible consequence of this increased formation of hydrogen bonds between polar side chains, as revealed by MD simulations (**Figure 2D**). Surficial residues on transmembrane (TM) helices (e.g., TM9/TM11, and TM5/TM7) show increased flexibility and form non-native polar contacts under detergent-free conditions, indicating that regions lacking screening are more inclined towards conformational heterogeneity.

Overall, **SE scores** and MD simulations consistently substantiate our observation that detergent removal leads to structural deterioration in vacuo. Such distortion are likely to render the protein susceptible to further disruption during ice-growth(*52*). When the protein is embedded in an amorphous ice film, counteracting the internal van-der-Waals forces that tend to self-compress the protein as the surrounding isotropic medium of similar density is restored(*38*). The growth temperature of 115 K was chosen to be close yet below crystalline-amorphous phase transition to allow for mobility of the water molecules and thus promote the formation of a smooth and flat, amorphous ice layer and structural relaxation at the protein surface.(*53, 54*) At the same time, complexes with only a few detergent molecules bound still degrade, likely because partial rehydration of exposed hydrophobic residues by water destabilises the protein (AmtB/AqpZ+15 DDM, see **Figure 1C** and **S3**). By contrast, if in the gas phase a larger amount of detergent are retained, the TM region is stabilised upon growth of the ice layer and improved resolution is found after embedding (see **Figure S7**). In addition, a large contingent of attached detergent may soften the impact upon landing, where soft non-covalent interactions absorb the majority of the collision energy avoiding rearrangement of the protein conformation.(*39, 50, 55, 56*)

### ESIBD maintains overall native membrane protein structures

As we are able to maintain sufficient detergent for structural preservation, we went further with our cryoEM data processing for two types of AmtB samples under maximal detergent retention: one with amorphous ice and one without (**Figure S8**). Both samples showed high-resolution information with identifiable secondary structure elements in 2D class averages, but the ice-free sample failed to yield a high-resolution 3D reconstruction. Instead, a low-resolution envelope is evident with limited internal features and a diffuse peripheral shell (**Figure S8**). This is consistent with previous reports from ice-free samples(*30, 37, 38*), indicating heterogeneity from dehydration or increased movement during radiation damage(*57*). Thus, gentle ion activation with maximal detergent retention, followed by amorphous ice growth, was used for obtaining 3D structure. This condition yielded the best preservation of the native AmtB structure.

We applied this approach to additional MP complexes spanning diverse oligomeric states, secondary structures, and domain compositions: AqpZ (tetramer, α-helical), ELIC (mixed α-helical and β-sheets with large soluble and TM domains), and OmpF (trimer, β-barrels) comparing each compared to a plunge-frozen control (**Figure S9**). Identity of the protein and their oligomeric states were confirmed by native MS using harsh activation conditions (**Figure 3A–C, i**), followed by deposition under gentle activation to preserve structural integrity (**Figure 3A–C, ii**.). Like AmtB, all membrane proteins produced 2D class averages very similar to plunge-frozen controls (**Figure 3A-C, iii**.) with characteristic α-helical and β-barrel features consistent with their architectures (**Figure 3A-D, iv**.). Notably, none of the ESIBD datasets exhibits visible density attributable to a detergent micelle, whereas plunge-frozen samples clearly show features consistent with a micelle.

**Fig. 3.**
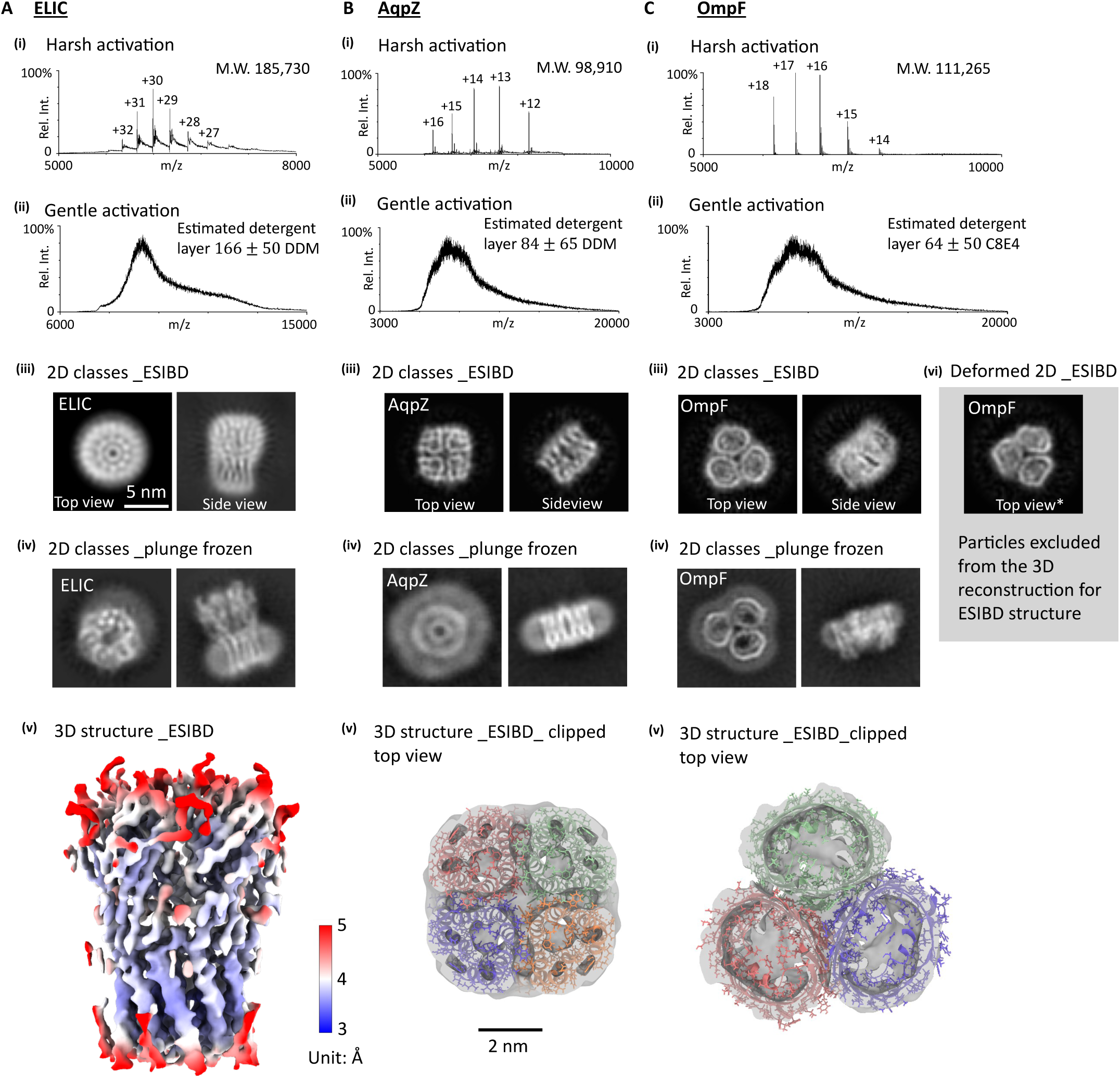
Mass spectra, cryoEM 2D classes and 3D density of membrane proteins (A) ELIC, (B) AqpZ, and (C) OmpF showing that structural integrity is preserved throughout ESIBD. Mass spectra of ELIC, AqpZ, and OmpF, following native ESI with **(i)** harsh and **(ii)** gentle activation conditions. The estimated detergent layer is obtained using the method shown in Figure S2, with parameters listed in Table S2. Representative 2D class averages after **(iii)** ESIBD and **(iv)** plunge freezing. Scale bars, 5 nm. **(v)** 3D reconstructions reveal the global architecture and symmetry of all membrane proteins. For ELIC, the density is coloured by local resolution; for AqpZ and OmpF, top-view clipped maps are shown overlaid on the models were fitted using the corresponding plunge-frozen maps. Scale bars, 2 nm. **(vi)** Deformed OmpF 2D class average (labelled *) observed in the ESIBD sample but excluded from the ESIBD 3D reconstruction.

All four proteins yield 3D reconstructions after gas-phase transfer, dehydration, and ice embedding (**Figure S10**), with the overall structure in good agreement with their native, plunge-frozen counterpart **(Figure S9)**. With a resolution below 3Å and well-resolved TM helices, the structure of AmtB reached the highest resolution and will be discussed in detail in the following section (**Figure 4A**). **AqpZ** exhibits density consistent with identifiable tetramer and TM helices (**Figure 3B**). **OmpF** retained the shape of its β-barrel TM domain, (**Figure 3C**), clearly visible in both, 2D classes and 3D density, albeit at a resolution which does not reveal the features of the secondary structure. The largest and most complex architecture, ELIC, with both soluble and TM domains, shows densities in which α-helical and β-sheet elements are clearly recognisable at better than 4 Å, with a global unmasked resolution of 5 Å (**Figure 3A**).

**Fig. 4.**
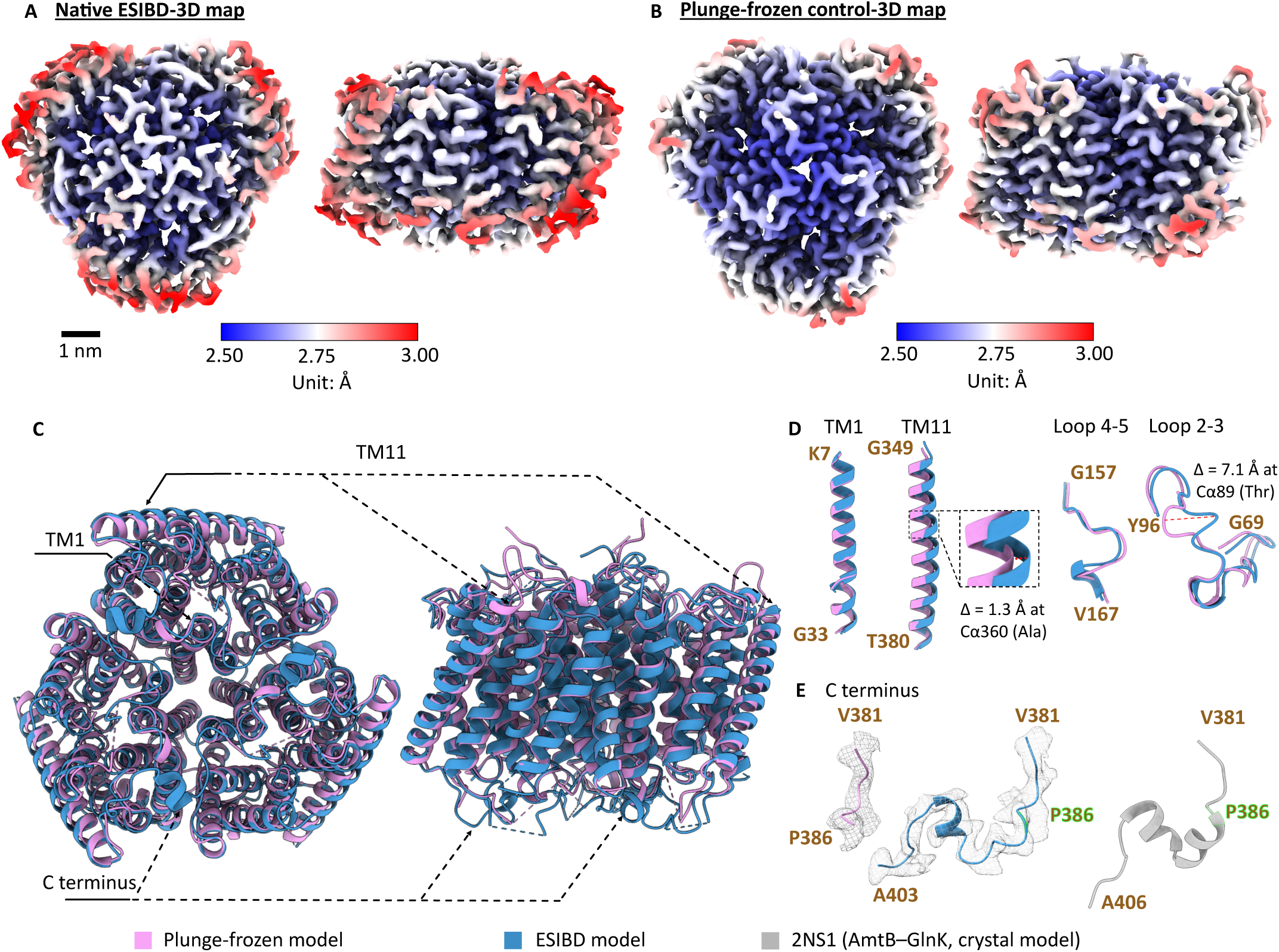
Comparison of AmtB ESIBD and plunge-frozen structures. **(A, B)** High-resolution cryoEM density maps of AmtB from native ESIBD (244,653 particles) and plunge-frozen control (239,700 particles) (details are in **Figure. S18**, **S19**). Scale bars, 1 nm. Color: local resolution. **(C)** Structural overlay of the full-length atomic models, plunge-frozen (pink) and ESIBD (blue) models. Both are generated by flexible fitting of PDB 1U77 into the 3.2 Å and 3.5 Å maps (unmasked resolution). **(D)** Examples of well-aligned motifs: TM1 and loop 4–5, and displaced motifs TM11 and loop 2–3 in the ESIBD model. **(E)** C-terminal segments across models, blue: ESIBD model, pink: plunge-frozen model, grey: 2NS1(*59*) which is the crystal model of AmtB complexed with the signal transduction protein GlnK. P386 is shown in fluorescent green in the ESIBD and 2NS1 models, indicating the truncation point of the plunge-frozen model.

Although overall architectures and folds are conserved across all membrane proteins, structural preservation and reconstruction resolution vary according to the complex. Comparing *in-vacuo* MD simulations and cryoEM data reveals dehydration-sensitive regions of the architecture linked to the observed structural changes. For example, the large β-barrel pore of OmpF undergoes a contraction, directly observed as dented barrels in some 2D-class averages (**Figure 3C top view\***). The MD simulations of OmpF in vacuum show a similarly contracted barrel, and high local RMSD at the solvent-exposed extracellular loops where deformation occurs (**Figure S11**). Here, hydrophilic regions inside the barrel become dehydrated and rearrange to compensate for missing polar interactions with water. AqpZ retains its tetrameric architecture, with the characteristic arrangement of α-helices evident in 2D class averages and appearing as blurred, tube-like densities in the 3D reconstruction, indicating that the overall fold of each subunit is preserved (**Figure 3B**). However, the low-resolution points to widespread heterogeneity acquired due to dehydration **(Figure S12).** MD in vacuum shows a significant contraction of the large, central, flexible and hydrophilic cavity at the periplasmic surface (**Figure S12**) which causes small rearrangements in the protein which can lead to heterogeneity.

ELIC is an ion channel protein with a central pore passing through the TM and extensive soluble domain. We were able to obtain a sub 4 Å reconstruction map (Bfactor = 220.1) in which features of α-helices and β-sheets can be resolved (**Figure 3A, S13A**). It is also almost fully resolved (with the exception of a short, flexible helix that is also missing in the PF-control), whereas several peripheral β-sheets and loops are absent in the soluble domain. Local-resolution in the TM domain is higher than in the soluble domain, a contrast not apparent in the plunge-frozen control (**Figure S13B**). Consistent with this observation, MD simulations match these observations, again suggesting how detergent adducts shield membrane-embedded regions and mitigate dehydration effects (**Figure S13C, D**).

The varying degrees of structural preservation across the different proteins can be described based on common principles. Dehydration removes bulk solvents, leading to unscreened interactions, predominantly in exposed hydrophilic patches. Dehydration-sensitive structural features share common properties, including mechanical flexibility and high solvent-exposure (*38*) due to topology or from nearby, solvent-filled cavities. Due to their reliance on hydration for stability they will undergo perturbations upon vacuum transfer in ESI. The effect of these perturbations on the cryoEM imaging depends on whether structural changes are coherent or random(*38*). Coherent changes preserve interpretable density despite structural change, whereas random changes, even if only small, lead to a loss in resolution, as in AqpZ. Retained detergent molecules can mitigate dehydration effects by screening otherwise exposed regions and providing polar binding partners via their head groups. This protection is effective for TM domains, as demonstrated by ELIC where the detergent-embedded TM region reaches higher resolution than the solvent-exposed extracellular domain.

### AmtB structure is retained in vacuum

For AmtB, where the compact architecture minimizes solvent-dependent features, detergent protection combined with ice embedding preserves the near-native structure at sub-3 Å resolution. In vacuum, AmtB is stabilised by a minimal detergent shell of about 60 ± 50 DDM molecules which corresponds to approximately a molecular monolayer coverage on the protein surface(schematic in **Figure S5**, one possible arrangement). We obtained a high-resolution 3D reconstruction of sub 3 Å (2.5 Å with masking in CryoSPARC, 3.5 Å without masking) which exhibited well-resolved helical structure, as shown in **Figure 4A**. The resolution we achieve is comparable to our plunge-frozen control (2.5 Å with masking in CryoSPARC, 3.2 Å without masking) with similar particle numbers (**Figure 4B**). The map captures the characteristic trimeric architecture of AmtB, including its ammonium-transport cavity, with clear density for TM helices, loops, and side chains throughout the complex, sufficient to enable confident modelling.

Strikingly, the vast majority (∼ 96%) of TM and surface-exposed regions are nearly identical between ESIBD and plunge-frozen maps, with an RMSD of 0.7 Å for 341 pruned atom pairs (1.5 Å across all 356 pairs) between the ESIBD and plunge frozen models, indicating near-complete preservation of the native structure. Helices and loops with continuous backbone density and clearly resolved side chains in ESIBD reconstructions are nearly identical to the plunge-frozen reference (**Figure S14, S15**), validating the native ESIBD approach to preserving MP structures in vacuum environment.

Despite the absence of a cryoEM-visible micelle in the ESIBD sample, the core TM structure of AmtB remains highly ordered, while peripheral helices and solvent-exposed loops show varying degrees of displacement. Structural alignment of the plunge-frozen and ESIBD models (**Figure 4C**) reveal minimal backbone deviations in the central core, with TM1 differing by 0.4 Å (measured at the Cα of residue 17) in local measurements (**Figure 4D**, left). In contrast, peripheral helices such as TM11 are displaced outward in the ESIBD structure by 1.3 Å (measured at the Cα of residue 360), likely due to the absence of the detergent micelle and the consequent loss of lateral packing forces, i.e., the surface tension of the micelle in contact with water (*7, 58*). This observation agrees well with our MD simulations (**Figure 2D**), where peripheral TM helix 11 exhibited strong distortions (per-helix localization details are shown in **Figure S16)**. As this displacement happens coherently for all proteins in the sample it has almost no effect on the local resolution of the final 3D reconstruction.

The loop regions of AmtB show different behaviours in ESIBD compared to the plunge-frozen control (**Figure 4D)**. Loop 4–5 remains consistent across both structural methods, suggesting rigidity, possibly due to its compact positioning near the helices. In contrast, loop 2–3 exhibits a pronounced conformational shift in the ESIBD model, with a backbone displacement of up to 7.1 Å relative to the plunge-frozen model, suggesting a coherent displacement to a new local energy minimum in the absence of solvent stabilization. Notably, the ESIBD map presents a longer C-terminal segment (V381–E403) compared with the plunge-frozen control, which only extends from V381–P386 (**Figure 4E**). **Intriguingly, it seems that** the vacuum environment stabilises the C-terminal segment, by strengthening intra-backbone hydrogen bonds and polar contacts, that is partly disordered in solution. The C terminus structure observed in the ESIBD map is remarkably similar to the conformation observed in a complex of *E. coli* AmtB–GlnK (PDB 2NS1) (*59*) and in AlphaFold3 model (**Figure S17**), suggesting that the local environment sampled during ESIBD has replicated one of the conformations that can be adopted by this dynamic region, potentially preserving a biologically relevant structure.

### Conclusions

By integrating native ESIBD with cryoEM, we determined structures of membrane proteins after native ESI, gas-phase transfer, soft-landing, and ice-embedding. Multiple protein architectures, α-helical and β-barrel, membrane and soluble domains, were shown to preserve their overall structures, demonstrating feasibility across diverse membrane proteins. Notably we obtained the structure of AmtB at sub-3 Å resolution, the first near-atomic structure of a membrane protein after transition through vacuum.

Our results reveal that detergent retention upon vacuum transfer of at least a molecular monolayer is sufficient and necessary for structural preservation. Amorphous ice embedding partially restores the hydration environment and radiation damage which can improve the resolution of structures from cryoEM, but only when the protein is sufficiently protected by detergent. In addition to stabilisation of hydrophobic regions, detergents can shield solvent-exposed regions during dehydration and prevent non-native polar contacts. However, structural preservation is architecture-dependent: solvent-sensitive structures are dynamic parts of the protein and features such as pores or channels will be perturbed due to the change in the potential energy landscape upon dehydration. Strikingly, we also found flexible regions of the protein, like the C terminus of AmtB, become more ordered than in solution due to stabilization by additional polar contacts.

The capability to soft-land, mass-select, and resolve membrane proteins at near-atomic resolution represents a significant advancement in gas-phase structural biology. Building on earlier studies showing that detergent micelles preserve protein structure during solution-to-gas transfer (*19*), our atomic-resolution structures provide a structural basis for interpreting native MS data with confidence. The direct MS-cryoEM pipeline enables linking structural and chemical characterisation of ligand binding, post-translational modifications, and heterogeneous complexes, areas where native MS excels.(*25, 60*) Currently, the method is well suited to compact, well-structured proteins rather than highly solvent-dependent architectures, but even for proteins showing dehydration-induced perturbation, the overall folds remain interpretable. The finding that adducts like detergents can stabilize protein structure upon dehydration, together with the possibility to integrate the nativeMS-ESIBD-cryoEM imaging pipeline with other methods such as laser flash melting (*61*) for annealing protein structure embedded in ice, establishes a foundation for future practical applications of gas-phase structural biology of soluble and membrane protein assemblies.

## Acknowledgments

We acknowledge support from Thermo Fisher Scientific who provided the Q Exactive UHMR mass spectrometer within the framework of a technology alliance partnership and the Aquilos sample transfer system. We would like to thank Melissa N. Webby, Abigail Ormrod and Colin Kleanthous (University of Oxford) for providing purified OmpF. We would like to thank Jani R. Bolla, Andrew Dolan, Benjamin F.Cooper, Haigang Song, Carla Kirschbaum, Tarick J El-Baba, Di Wu, Benjamin Mallada, and Alejandro Lynch Gonzalez for helpful discussions and suggestions. We also would like to acknowledge support from the COSMIC microscope facility. Computations were performed at Advanced Research Computing resources in the University of Oxford and NSC Tetralith provided by the National Academic Infrastructure for Supercomputing in Sweden (NAISS), partially funded by the Swedish Research Council through grant agreement no. 2022-06725, and Oxford ARC supercomputing cluster in the UK.

## Funding

UK Research and Innovation (UKRI) Horizon Europe Guarantee: Marie Skłodowska-Curie Actions (MSCA) Postdoctoral Fellowship EP/Z001684/1(J.F.)

Junior Research Fellowship from Wolfson College in University of Oxford (J.F.)

Wellcome Trust Grant 228310/Z/23/Z (C.D.)

Swedish Research Council through Grant Agreement No. 2020-04825 (E.G.M., L.J.P.)

C.F. Liljewalchs travel stipend (L.J.P.)

Wellcome Trust Grant 218482/Z/19/Z (L.T.S.)

UK Research and Innovation (UKRI) Medical Research Council (MRC) MR/V028839/1 (C.V.R.)

Wellcome Trust grant number 221795/Z/20/Z (C.V.R.)

UK Research and Innovation (UKRI) Engineering and Physical Sciences Research Council (EPSRC) EP/V051474/1 (S.R.)

UK Research and Innovation (UKRI) Biotechnology and Biological Sciences Research Council (BBSRC) BB/V019694/1 (S.R.)

## Author contributions

Conceptualization: JF, SR, CVR, LB

Hardware of the instrument: TE, LE, SR

Native MS experiment: JF, LE, TE

ESIBD experiment, imaging: JF, LE, TE

ESIBD sample cryoEM data analysis: JF, CD, LE, CvH, LTS, SBK, LB

Atomic model building: JF, CD, LTS, LB

MD simulation and solvent exposure calculation: CvH, LJP, LE, NN, EGM

Plunge-frozen control experiment and data analysis: CD, LB

Protein purification: AOO, AQ, SM, JF, NK

Interpretation of the results: all the authors

Supervision: SR, CVR, LB

Writing – original draft: JF, SR

Writing – review & editing: all the authors

## Data and materials availability

Cryo-EM density maps and atomic models will be deposited in the Electron Microscopy Data Bank (EMDB) and the Protein Data Bank (PDB), respectively, and will be released upon publication. The MD data will be deposited to a Zenodo repository. All data are available in the main text or the supplementary materials.

## Reference

1. G. Von Heijne, Membrane-protein topology. Nature reviews Molecular cell biology 7, 909–918 (2006).

2. J. P. Overington, B. Al-Lazikani, A. L. Hopkins, How many drug targets are there? Nature reviews Drug discovery 5, 993–996 (2006).

3. A. Laganowsky et al., Membrane proteins bind lipids selectively to modulate their structure and function. Nature 510, 172–175 (2014).

4. S. A. Lawrence, A. Dolan, M. M. Miller, C. V. Robinson, Membrane Protein Complexity Revealed Through Native Mass Spectrometry. Annual Review of Biochemistry 94, (2025).

5. D. S. Chorev et al., Protein assemblies ejected directly from native membranes yield complexes for mass spectrometry. Science 362, 829–834 (2018).

6. S. M. Fantin et al., Collision induced unfolding classifies ligands bound to the integral membrane translocator protein. Analytical chemistry 91, 15469–15476 (2019).

7. S. C. Wang et al., Ion mobility mass spectrometry of two tetrameric membrane protein complexes reveals compact structures and differences in stability and packing. Journal of the American Chemical Society 132, 15468–15470 (2010).

8. E. Christofi, P. Barran, Ion mobility mass spectrometry (IM-MS) for structural biology: insights gained by measuring mass, charge, and collision cross section. Chemical reviews 123, 2902–2949 (2023).

9. C. Martens, M. Shekhar, A. M. Lau, E. Tajkhorshid, A. Politis, Integrating hydrogen–deuterium exchange mass spectrometry with molecular dynamics simulations to probe lipid-modulated conformational changes in membrane proteins. Nature protocols 14, 3183–3204 (2019).

10. X. Qiu et al., Coupling and Activation of the β1 Adrenergic Receptor-The Role of the Third Intracellular Loop. Journal of the American Chemical Society 146, 28527–28537 (2024).

11. R. Friemann, D. S. Larsson, Y. Wang, D. van der Spoel, Molecular dynamics simulations of a membrane protein− micelle complex in vacuo. Journal of the American Chemical Society 131, 16606–16607 (2009).

12. M. Landreh et al., Integrating mass spectrometry with MD simulations reveals the role of lipids in Na+/H+ antiporters. Nature communications 8, 13993 (2017).

13. M. T. Agasid, C. V. Robinson, Probing membrane protein–lipid interactions. Current Opinion in Structural Biology 69, 78–85 (2021).

14. J. Marcoux, C. V. Robinson, Twenty years of gas phase structural biology. Structure 21, 1541–1550 (2013).

15. S. Kumar et al., Native Mass Spectrometry of Membrane Protein–Lipid Interactions in Different Detergent Environments. Analytical Chemistry 96, 16768–16776 (2024).

16. P. G. Wolynes, Biomolecular folding in vacuo!!!(?). Proceedings of the National Academy of Sciences 92, 2426–2427 (1995).

17. M. F. Jarrold, Unfolding, refolding, and hydration of proteins in the gas phase. Accounts of Chemical Research 32, 360–367 (1999).

18. J. L. Benesch, C. V. Robinson, Dehydrated but unharmed. Nature 462, 576–577 (2009).

19. N. P. Barrera, N. Di Bartolo, P. J. Booth, C. V. Robinson, Micelles protect membrane complexes from solution to vacuum. Science 321, 243–246 (2008).

20. A. C. Leney, A. J. Heck, Native mass spectrometry: what is in the name? Journal of the American Society for Mass Spectrometry 28, 5–13 (2016).

21. R. M. Bill et al., Overcoming barriers to membrane protein structure determination. Nature biotechnology 29, 335–340 (2011).

22. H. E. Autzen, D. Julius, Y. Cheng, Membrane mimetic systems in CryoEM: keeping membrane proteins in their native environment. Current opinion in structural biology 58, 259–268 (2019).

23. B. C. Choy, R. J. Cater, F. Mancia, E. E. Pryor Jr, A 10-year meta-analysis of membrane protein structural biology: Detergents, membrane mimetics, and structure determination techniques. Biochimica et Biophysica Acta (BBA)-Biomembranes 1863, 183533 (2021).

24. D. Hardy, R. M. Bill, A. Jawhari, A. J. Rothnie, Overcoming bottlenecks in the membrane protein structural biology pipeline. Biochemical Society Transactions 44, 838–844 (2016).

25. L. Kuhlen et al., Structure of the core of the type III secretion system export apparatus. Nature structural & molecular biology 25, 583–590 (2018).

26. S. Rauschenbach, M. Ternes, L. Harnau, K. Kern, Mass spectrometry as a preparative tool for the surface science of large molecules. Annual Review of Analytical Chemistry 9, 473–498 (2016).

27. J. Seibel, K. Anggara, M. Delbianco, S. Rauschenbach, Scanning Probe Microscopy Characterization of Biomolecules enabled by Mass - Selective, Soft - landing Electrospray Ion Beam Deposition. ChemPhysChem 25, e202400419 (2024).

28. G. E. Johnson, Q. Hu, J. Laskin, Soft landing of complex molecules on surfaces. Annual Review of Analytical Chemistry 4, 83–104 (2011).

29. B. Gologan, J. R. Green, J. Alvarez, J. Laskin, R. G. Cooks, Ion/surface reactions and ion soft-landing. Physical Chemistry Chemical Physics 7, 1490–1500 (2005).

30. T. K. Esser et al., Cryo-EM samples of gas-phase purified protein assemblies using native electrospray ion-beam deposition. Faraday discussions 240, 67–80 (2022).

31. M. S. Westphall et al., Three-dimensional structure determination of protein complexes using matrix-landing mass spectrometry. Nature communications 13, 2276 (2022).

32. H. Ochner, S. Rauschenbach, L. Malavolti, Electrospray ion beam deposition plus low-energy electron holography as a tool for imaging individual biomolecules. Essays in Biochemistry 67, 151– 163 (2023).

33. Z. Ouyang et al., Preparing protein microarrays by soft-landing of mass-selected ions. Science 301, 1351–1354 (2003).

34. G. Siuzdak et al., Mass spectrometry and viral analysis. Chemistry & biology 3, 45–48 (1996).

35. V. A. Mikhailov, T. H. Mize, J. L. Benesch, C. V. Robinson, Mass-selective soft-landing of protein assemblies with controlled landing energies. Analytical chemistry 86, 8321–8328 (2014).

36. Z. Deng et al., A close look at proteins: submolecular resolution of two-and three-dimensionally folded cytochrome c at surfaces. Nano letters 12, 2452–2458 (2012).

37. T. K. Esser et al., Cryo-EM of soft-landed β-galactosidase: Gas-phase and native structures are remarkably similar. Science advances 10, eadl4628 (2024).

38. L. Eriksson, et al., High-resolution cryoEM structure determination of soluble proteins after soft-landing ESIBD. *arXiv preprint arXiv:2503.22364*, (2025).

39. E. Reading et al., The role of the detergent micelle in preserving the structure of membrane proteins in the gas phase. Angewandte Chemie International Edition 54, 4577–4581 (2015).

40. S. Tamara, M. A. den Boer, A. J. Heck, High-resolution native mass spectrometry. Chemical reviews 122, 7269–7326 (2021).

41. P. Fremdling et al., A preparative mass spectrometer to deposit intact large native protein complexes. ACS nano 16, 14443–14455 (2022).

42. J. W. Patrick et al., Allostery revealed within lipid binding events to membrane proteins. Proceedings of the National Academy of Sciences 115, 2976–2981 (2018).

43. S. M. Dixit, D. A. Polasky, B. T. Ruotolo, Collision induced unfolding of isolated proteins in the gas phase: past, present, and future. Current opinion in chemical biology 42, 93–100 (2018).

44. C. Eldrid et al., Cyclic ion mobility–collision activation experiments elucidate protein behavior in the gas phase. Journal of the American Society for Mass Spectrometry 32, 1545–1552 (2021).

45. J. S. Brodbelt, Ion activation methods for peptides and proteins. Analytical chemistry 88, 30–51 (2016).

46. B. Lee, F. M. Richards, The interpretation of protein structures: estimation of static accessibility. Journal of molecular biology 55, 379–IN374 (1971).

47. A. Shrake, J. A. Rupley, Environment and exposure to solvent of protein atoms. Lysozyme and insulin. Journal of molecular biology 79, 351–371 (1973).

48. S. L. Rouse, J. Marcoux, C. V. Robinson, M. S. Sansom, Dodecyl maltoside protects membrane proteins in vacuo. Biophysical journal 105, 648–656 (2013).

49. D. van der Spoel, E. G. Marklund, D. S. Larsson, C. Caleman, Proteins, lipids, and water in the gas phase. Macromolecular bioscience 11, 50–59 (2011).

50. K. Breuker, F. W. McLafferty, Stepwise evolution of protein native structure with electrospray into the gas phase, 10− 12 to 102 s. Proceedings of the National Academy of Sciences 105, 18145–18152 (2008).

51. Y. Zhang et al., Hydrophobic Shielding Preserves Transmembrane Secondary Structure in the Gas Phase. The Journal of Physical Chemistry Letters 16, 12875–12881 (2025).

52. P. H. Handle, T. Loerting, F. Sciortino, Supercooled and glassy water: Metastable liquid (s), amorphous solid (s), and a no-man’s land. Proceedings of the National Academy of Sciences 114, 13336–13344 (2017).

53. K. Amann-Winkel et al., Water’s second glass transition. Proceedings of the National Academy of Sciences 110, 17720–17725 (2013).

54. R. Buchner, J. Barthel, J. Stauber, The dielectric relaxation of water between 0 C and 35 C. Chemical Physics Letters 306, 57–63 (1999).

55. K. Anggara et al., Landing proteins on graphene trampoline preserves their gas-phase folding on the surface. ACS Central Science 9, 151–158 (2022).

56. K. Anggara et al., Exploring the molecular conformation space by soft molecule–surface collision. Journal of the American chemical society 142, 21420–21427 (2020).

57. M. J. Peet, R. Henderson, C. J. Russo, The energy dependence of contrast and damage in electron cryomicroscopy of biological molecules. Ultramicroscopy 203, 125–131 (2019).

58. C. Chipot et al., Perturbations of native membrane protein structure in alkyl phosphocholine detergents: a critical assessment of NMR and biophysical studies. Chemical reviews 118, 3559– 3607 (2018).

59. F. Gruswitz, J. O’Connell III, R. M. Stroud, Inhibitory complex of the transmembrane ammonia channel, AmtB, and the cytosolic regulatory protein, GlnK, at 1.96 Å. Proceedings of the National Academy of Sciences 104, 42–47 (2007).

60. H.-Y. Yen et al., Ligand binding to a G protein–coupled receptor captured in a mass spectrometer. Science advances 3, e1701016 (2017).

61. S. V. Barrass et al., Cryo-EM Sample Preparation with Soft-Landing and Laser Flash Melting. bioRxiv, 2025.2006. 2005.657968 (2025).

