## Supplementary Material for "Membrane proteins retain native architecture through native ESI and soft-landing"

**The PDF file includes:**

Materials and methods

Figs. S1 to S25

Tables S1 to S6

Reference

#### Materials and methods

##### Protein preparation

Membrane proteins were expressed and purified as previously described (*1-4*). Briefly, AmtB, AqpZ, and ELIC were expressed in *E. coli* host. Membrane fraction was solubilized in 1-2% n-dodecyl-β-D-maltoside (DDM, $m_w$=510.62 g/mol). All purification buffers contained DDM at 2× the critical micelle concentration (CMC), ensuring adequate micelle encapsulation during purification. OmpF, a β-barrel outer membrane protein, was purified in n-octyl-β-D-glucoside (β-OG) at 2× CMC, which is better suited for stabilizing its structure(*5*).

##### Native MS

Nano-electrospray ionization (nESI) emitters were produced in-house from borosilicate glass capillaries (30-0042, Harvard Bioscience) using a P-1000 pipette puller (Sutter Instrument) and subsequently coated with gold via sputtering (108A/SE, Cressington). Each emitter was filled with up to 5 µL of protein solution, and the tip was carefully trimmed to produce an aperture ranging from 0.1 to 10 µm(*6*). Electrospray was initiated by applying a voltage of 1.2 kV, with a gentle backing pressure of less than 100 mbar above atmospheric level. Transfer capillary temperature was maintained at 150°C for AmtB, AqpZ, OmpF and 50°C for ELIC. For better spray, proteins were desalted twice using Zeba micro spin desalting columns (7K MWCO), equilibrated with 40µL of 200 mM ammonium acetate(pH7.0) containing 2× CMC of detergent. The final protein concentration used was 10 µM. AmtB and AqpZ were introduced into the mass spectrometer in solutions containing DDM micelles with C8E4 detergent present. Only DDM adducts were identified in the mass spectra and thus explicitly considered further analysis. We note that the presence of residual or weakly associated C8E4 molecules cannot be excluded, particularly in the gentle-activation conditions (Figure. S2 and S3). OmpF was introduced into the mass spectrometer from C8E4 micelles and we observe C8E4 adducts in the mass spectra. ELIC was sprayed from DDM micelles.

**ESIBD workflow**

We used a modified Thermo Scientific Q Exactive UHMR mass spectrometer, extended by a custom deposition stage for native ESIBD (Fig. S1). In addition to high-resolution native MS, our instrument features: a collision cell for ion activation or thermalization, guiding ion optics, a cryogenically cooled deposition stage, and a cryo-shuttle. The method introduced before (*7-9*) is schematically shown in Figure S1. Before deposition, ion beam parameters, including ion activation trapping voltage, mass selection, beam intensity, and deposition energy, are optimized to ensure controlled landing of protein ions. During ion transfer, the effects of different numbers of detergent molecules surrounding membrane proteins can be studied by modulating the ion transfer optics during activation.

The deposition stage, kept at ultrahigh vacuum conditions ($P < 10^{-8}$ mbar) maintains a consistent temperature of the TEM grid at 115 K (*8*). The membrane protein ions land individually onto a cold carbon film grid used for CryoEM, ensuring immobilization on the surface.(*10*) To avoid radiation damage in EM(*11*) and enable hydrogen bonding and van-der-Waals interactions which are crucial for structural integrity, we grow a thin layer of amorphous ice over the landed proteins. We temporarily increase the partial pressure of water in the chamber to $5\times10^{-5}$ mbar while keeping the shuttle inside the deposition stage at 115K as shown in Fig. S1. The ice growth temperature of 115 K was chosen to be close yet below the crystalline-amorphous phase transition to allow for mobility of water molecules. (*12*) Thus these conditions promote both the formation of a homogenous, smooth and flat, 20 nm film of low-density amorphous ice layer and structural relaxation at the protein surface.(*8, 12, 13*) Once the deposition and ice growth are complete, the cryo-shuttle containing the shielded grid is transferred from the vacuum chamber into liquid nitrogen via a dry nitrogen-purged transfer system, then transferred under liquid nitrogen to the microscope to avoid devitrification. By operating under soft-ionization, soft-landing and cryogenic conditions, we minimize thermal diffusion and structural rearrangements of the landed membrane protein- detergent complex, during the desolvation, deposition process and throughout subsequent sample handling.

**Preparation of the plunge-frozen control grids**

The control samples were prepared using Quantifoil grids of varying holes and mesh size. Grids were glow-discharged using the plasma cleaner. 3 μL of solution were applied to the grid, followed by blotting and plunging into liquid ethane, using a Vitrobot Mark IV plunger. The details for each protein can be found in Table S3.

**Preparation of the native ESIBD grid**

Grids were prepared and transferred as described in the main text. Figure S1 shows an overview of the deposition instrument and cryo deposition stage described in more detail elsewhere (*7, 8, 10, 14*). At the beginning of a deposition experiment, clipped TEM grids are loaded into the shuttle at room temperature. Here, copper TEM grids from Quantifoil with mesh size 400, R2/1 holey carbon covered with 2-nm amorphous carbon were used for AmtB and ELIC. Gold TEM grids from Quantifoil with mesh size 300, R1.2/1.3 holey carbon covered with home-made amorphous carbon (Emitech K950X Carbon Coater) were used for AqpZ and OmpF. The shuttle was loaded into the stage a few minutes before deposition to allow for thermalization to the stage, which was held at 115 K using a cryogenic cooler as described before.

The beam energy is measured using a retarding-grid energy detector. The deposited charge is monitored by a picoammeter to control protein coverage. Protein ions are deposited with a deposition energy of 1.5 eV per charge, by applying a corresponding potential to the TEM grids. Close to monolayer coverage in the grid center is typically achieved in 90 min, and lower densities for efficient data collection can be found further from the center if needed. After completion, transfer into liquid nitrogen is done in less than 2 min as described before.

**Movie acquisition and processing**

All cryoEM data were acquired on a Thermo Scientific Titan Krios transmission electron microscope operating at 300 kV and equipped with a BioQuantum energy filter (slit width: 20 eV) and a Gatan K3 direct electron detector. Data collection was performed at the COSMIC CryoEM facility, University of Oxford, using automated acquisition via EPU software (Thermo Scientific). Movies were recorded in TIFF format at a nominal magnification of 105,000×, corresponding to a calibrated pixel size of 0.83 or 0.85 Å (details can be found in SI table 4-6). Images were collected with a total electron dose of 40 $e^{-}/Å^{2}$ and a defocus range of −1.75 µm to −3 µm.

Data processing was carried out in cryoSPARC. Motion correction and contrast transfer function (CTF) estimation were performed using Patch Motion Correction and Patch CTF modules, respectively. Particles were initially picked manually to generate templates for template-based autopicking, then extracted in 360-pixel boxes for AmtB and OmpF, 448-pixel boxes for AqpZ, and 416-pixel boxes for ELIC. Subsequent processing included multiple rounds of 2D classification, followed by heterogeneous, homogeneous, non-uniform 3D refinement, and local refinement. Local resolution estimation and local filtering were performed to assess map quality.

Symmetrical constraints were applied during refinement: C3 for AmtB, C4 for AqpZ, C3 for OmpF, and C5 for ELIC. Final density maps and associated visualization figures and movies were generated using UCSF ChimeraX (*15*). Further acquisition and processing parameters are provided in Figures S17-24 and Table S4-6.

**Model building**

A previously published crystal model of AmtB (PDB:1U77) was used as a starting model for both ESIBD and plunge-frozen model building. All the water molecules were removed and all the MSE residues have been changed to MET. The model was aligned and fitted into the 3D ESIBD map and Plunge-frozen map in ChimeraX, and relaxed into the maps using ISOLDE(*15*), resulting in good agreement of secondary structure elements. The crystal structures fit the reconstructions well for most regions, except for both the N and C termini and the loops, which required refinement of their relative angles to the rest of the structure. A single chain model was manually fitted using COOT (*16*) with residues mutated as necessary to match the purified AmtB sequence. This was followed by real-space refinement using 1 to 5 cycles and 100 iterations to optimize the model-map fit and reduce clashes. The resulting models were iteratively rebuilt in COOT and then refined in PHENIX (*17*) until completion. A three-chain model in C3 symmetry was generated using the Map Symmetry and Apply NCS Operators jobs in PHENIX before real-space refinement as previously described with restraints of secondary structure and non-crystallographic symmetry (NCS). Previously published models of OmpF (PDB: 6WTZ, EM) and AqpZ (2ABM, XRC) were used as starting models for plunge-frozen model building. Model Angelo (*18*), available in the RELION v5 suite (*19*), was used to generate an ELIC starting model *de novo* for plunge-frozen model building. Similar operations as described for AmtB were completed for the plunge-frozen maps of OmpF, AqpZ and ELIC with C3, C4 and C5 symmetry applied respectively. Where required, map handedness was inverted in RELION v5 using the inverted hand function from Relion image handler before aligning or generating starting models.

**Molecular dynamics**

Simulations of AmtB, OmpF, ELIC, and AqpZ were initiated from published PDB structures (1U77, 6WTZ, 8F35, and 2ABM respectively). The PDB files were prepared for simulation using the CHARMM-GUI micelle builder (*20-22*) to produce a micelle composed of 150 detergent molecules. CHARMM-GUI was also used to predict protonation states at pH 7.0. Some acidic side chains were neutralized (OmpF: GLU296, ASP7 in subunit A, ELIC: ASP122, ASP13 in subunit A, AqpZ: GLU8, GLU31, GLU80, ASP110 in subunit A, AmtB: none). The systems were solvated using TIP3P water molecules and NaCl ions were added to a concentration of 150 mM to neutralise any net charge.

Each protein was simulated under three different conditions, reflecting the differing number of the detergent adducts observed when varying the in-source trapping voltage in native MS (150, 15, and 0 DDM detergent molecules, respectively). Three repeats were conducted for each number of adducts, resulting in a total of 9 simulations per protein system. All simulations were conducted using GROMACS 2023.4 (*23, 24*) on the Oxford ARC and NCS Tetralith supercomputing clusters.

For OmpF, Elic and AqpZ each of the 9 simulation repeats was first equilibrated in solution in a micelle containing 150 detergent molecules. For each repeat, a solvated, neutralized protein with 150 detergent adducts generated from CHARMM-GUI was subjected to energy minimisation using the steepest-descent algorithm, followed by NVT and NPT equilibration using the standard CHARMM-GUI protocol After minimization, a series of restrained NPT equilibration simulations totaling 1.875 ns was conducted. LINCS constraints were applied to all hydrogen bonds. The system was equilibrated to 310K (OmpF, Elic and AqpZ) or 300K (AmtB) and atmospheric pressure using the Bussi-Donadio-Parrinello ("v-rescale") thermostat (*25*) and "C-rescale" barostat (*26*). For simulations in vacuum, the centre-of-mass translational and rotational velocities were removed every 10th time step. Solvent, membrane and protein were temperature-coupled separately. During the equilibration, restraints on backbone, side chains, dihedrals and detergent molecules were gradually relaxed. Restraints of 4000 kJ $mol^{-1}$ $nm^{2}$, 2000 kJ $mol^{-1}$ $nm^{-2}$, 1000 kJ $mol^{-1}$ $nm^{-2}$ and 1000 kJ $mol^{-1}$ $rad^{-2}$ were applied to backbone, side chains, detergent, and dihedral angles respectively for the first 125 ps using 1-fs timesteps. This was followed by a further 125 ps with 2000 kJ $mol^{-1}$ $nm^{2}$, 1000 kJ $mol^{-1}$ $nm^{-2}$, 0 kJ $mol^{-1}$ $nm^{-2}$ and 400 kJ $mol^{-1}$ $rad^{-2}$ and another 125 ps with 1000 kJ $mol^{-1}$ $nm^{2}$, 500 kJ $mol^{-1}$ $nm^{-2}$, 0 kJ $mol^{-1}$ $nm^{-2}$ and 200 kJ $mol^{-1}$ $rad^{-2}$. The time step was then increased to 2 fs and the system simulated for 500 ps with restraints of 500 kJ $mol^{-1}$ $nm^{2}$, 200 kJ $mol^{-1}$ $nm^{-2}$, 0 kJ $mol^{-1}$ $nm^{-2}$ and 200 kJ $mol^{-1}$ $rad^{-2}$. Two additional 500-ps equilibrations were then conducted, relaxing the restrains first to 200 kJ $mol^{-1}$ $nm^{2}$, 50 kJ $mol^{-1}$ $nm^{-2}$, 0 kJ $mol^{-1}$ $nm^{-2}$ and 100 kJ $mol^{-1}$ $rad^{-2}$ and finally to 50 kJ $mol^{-1}$ $nm^{2}$, 0 kJ $mol^{-1}$ $nm^{-2}$, 0 kJ $mol^{-1}$ $nm^{-2}$ and 0 kJ $mol^{-1}$. Following NPT equilibration, each repeat was simulated in solution without restraints for 3 ns with a 2 fs timestep and then prepared for the transition to the gas-phase.

After equilibration with a micelle of 150 detergent adducts, each repeat was individually prepared for transfer into vacuum, with detergent being removed where necessary. To model charge additions during the ESI-process, the most solvent-accessible titratable residues (aspartate, glutamate, and the carboxyl group of the C-terminus) were identified using the double-cubic lattice method implemented in GROMACS and protonated until the principal charge state observed in the native MS was reached (OmpF = +17, ELIC = +15, AqpZ = +14). For vacuum simulations with only 15 detergent molecules, the 15 detergent molecules with the greatest number of contacts were retained, where contacts were defined as the number of protein atoms within 4.5 Å of a given detergent atom. For gas-phase simulations without any detergent, all detergent molecules were deleted during this stage. AmtB was also equilibrated using the default CHARMM-GUI protocol described above. After the final restrained equilibration, AmtB was simulated in solution for 10 ns and structures for simulations in vacuum with 150, 15 and 0 detergent adducts were extracted from the final 3 ns of the equilibration. Detergent was then removed and charges were added as described above (Amtb charge = +16). For each starting structure, 3 replicates were minimized and equilibrated in vacuum.

All systems were equilibrate in vacuum using the following procedure: after the deletion of all solvent molecules and ions, virtual sites for hydrogen were added to the protein and to detergent using MkVsites as implemented in GROMACS (*27*). To deal with long-range charge interactions in the gas phase, a pseudo-periodic boundary conditions (pseudo-PBC) approach was used to prevent interactions across PBCs, meaning that the system was placed in a 999.9 x 999.9 x 999.9 Å$^3$ simulation box (*28*) and the cutoff radii for Coulomb and van der Waals forces were set to 333.3 nm. Each system then underwent a further round of energy minimization via steepest-descent and 100 ps of NVT equilibration for protein and detergent, separately, using the Berendsen thermostat (*29*) (note that no barostat was used as the system was simulation under vacuum conditions). Production simulations lasting for 300 ns were conducted using the v-rescale thermostat at 300 K with a 5-fs timestep. Frames were written every 20 ps.

Trajectory analysis was performed using the MDAnalysis package (*30, 31*). Hydrogen bonds were measured using cutoffs of 3.5 Å and 150°. Helical bending was measured using the Bendix-plugin (*32*) in VMD with results exported and analyzed in python. Pore collapse was visualised and quantified with CHAP (Channel Annotation Package)(*33*).

We note that ion activation in MS involves more than the simple removal of water and detergent, as energetic collisions with background gas transfer additional energy to proteins. The timescales for energy redistribution depend on collision rates and energies and may lead to different structural change mechanisms. In the present simulations, we therefore isolate the effects of water and detergent removal without explicitly modeling protein–gas collisions.

**Solvent exposure definition and calculation**

To assess the protection effect of detergent molecules on membrane proteins, and also the stability differences of the membrane protein with different detergent protection, under gas-phase and dehydration conditions, we computed a simplified solvent exposure score from all-atom structural models. (*8*) Here, in the membrane protein system, solvent-exposure scores are defined as the ratio of solvent to protein/ detergent atoms within a 5 nm cutoff. This approach allows us to quantify how exposed each protein atom is to its surrounding environment, either other protein atoms or detergent molecules—based on pairwise spatial proximity. Proteins are displayed in ribbon view, and per-residue scores are obtained by averaging the atomic scores in each residue. Using a python script (https://github.com/lukasaerik/solvent_exposure), the solvent exposure score is calculated for the MD solvent-equilibrated models, including any DDM molecules. For each non-hydrogenic atom, $i$, we sum up $d_{ij}^{-2}$, where $d_{ij}$ is the distance (measured in Ångström) between the atom and all other non-hydrogenic atoms, $j$. Each atom's score is then subtracted from the maximal score obtained from all proteins considered. This is visualized in ChimeraX by coloring the protein by the calculated scores.

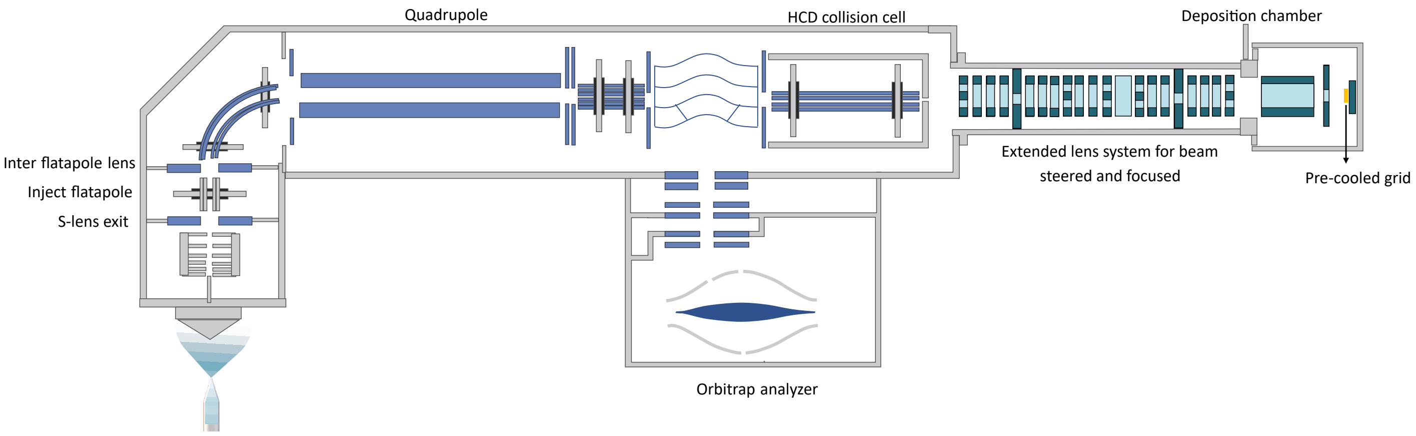


**Fig. S1.**

**Overview of the native electrospray ion beam deposition instrument.** Membrane proteins are ionized using a nano-electrospray ionization source and then transferred into the gas phase. After the ion activation, ion transfer, and mass selection process, the ion beam is deposited onto a pre-cooled grid in the deposition chamber. The ion beam is steered and focused in the extended lens system. Detailed designed can be found in previous work (*9*). This instrument is modified from a commercial mass spectrometer (Thermo Scientific Q Exactive UHMR instrument).

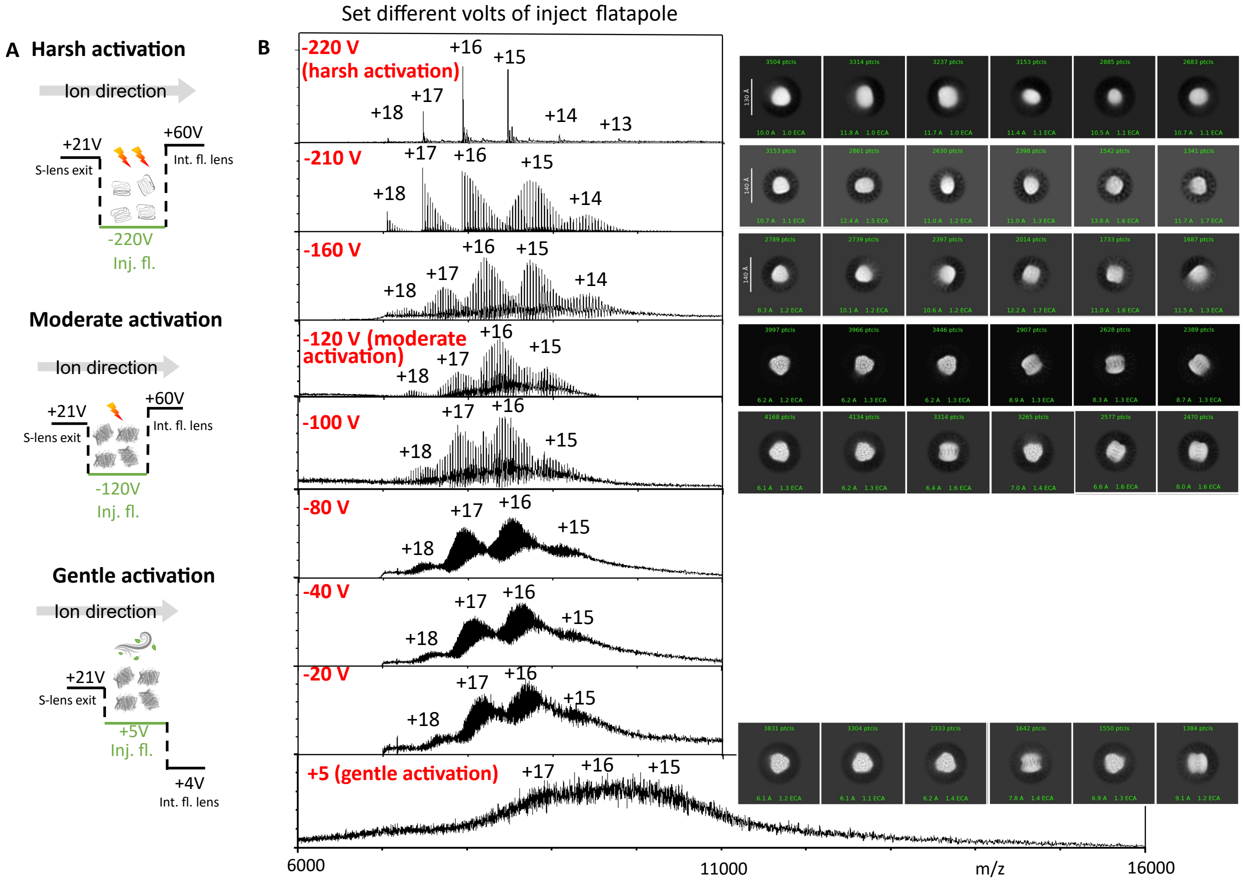


**Fig. S2.**

**Mass spectra and cryoEM of AmtB under different activation mode without ice.** In the ion activation mode, ions will be trapped for 10 ms within a potential well of variable depth in a chamber filled with N2 as collision gas at a pressure of around 2 mbar. **(A)** Schematic of different activation modes. 'Inj. fl.' means inject flatapole . 'Int. fl. lens' means inter -flatapole lens. **(B)** Charge state distribution of AmtB as a function of activation voltages and related cryoEM 2D class averages from datasets with comparable particle numbers . We assume that the charge state distribution doesn 't change during the gradient of activation voltages. Under this assumption, we can determine the charge state from other mass spectra and then calculate the number of bound detergent molecules. From the spectrum, we obtain $z_{main} = 16$ and $(m/z)_{main} = 9800$. The mass of AmtB is taken as $m_{AmtB} = 126.6\ kDa$ , and the mass of one DDM molecule is $m_{DDM} = 510\ Da$ and The number of associated DDM molecules, $N_{DDM}$, and its uncertainty, $\Delta N_{DDM}$, are estimated using:

$$N_{DDM} = ((m/z)_{main} \times z_{main} - m_{AmtB})\frac{1}{m_{DDM}}$$

giving $N_{DDM} \approx 60$. The uncertainty is estimated by error propagation:

$$\Delta N_{DDM} = \left|\frac{\partial N}{\partial z}\right|\Delta z + \left|\frac{\partial N}{\partial \frac{m}{z}}\right|\Delta\frac{m}{z}$$

Using $\Delta z = 1$, and $\Delta\frac{m}{z} \approx 1000$, we obtain $\Delta N_{DDM} \approx 50$. Thus, $N_{DDM} \pm \Delta N_{DDM} = 60 \pm 50$.

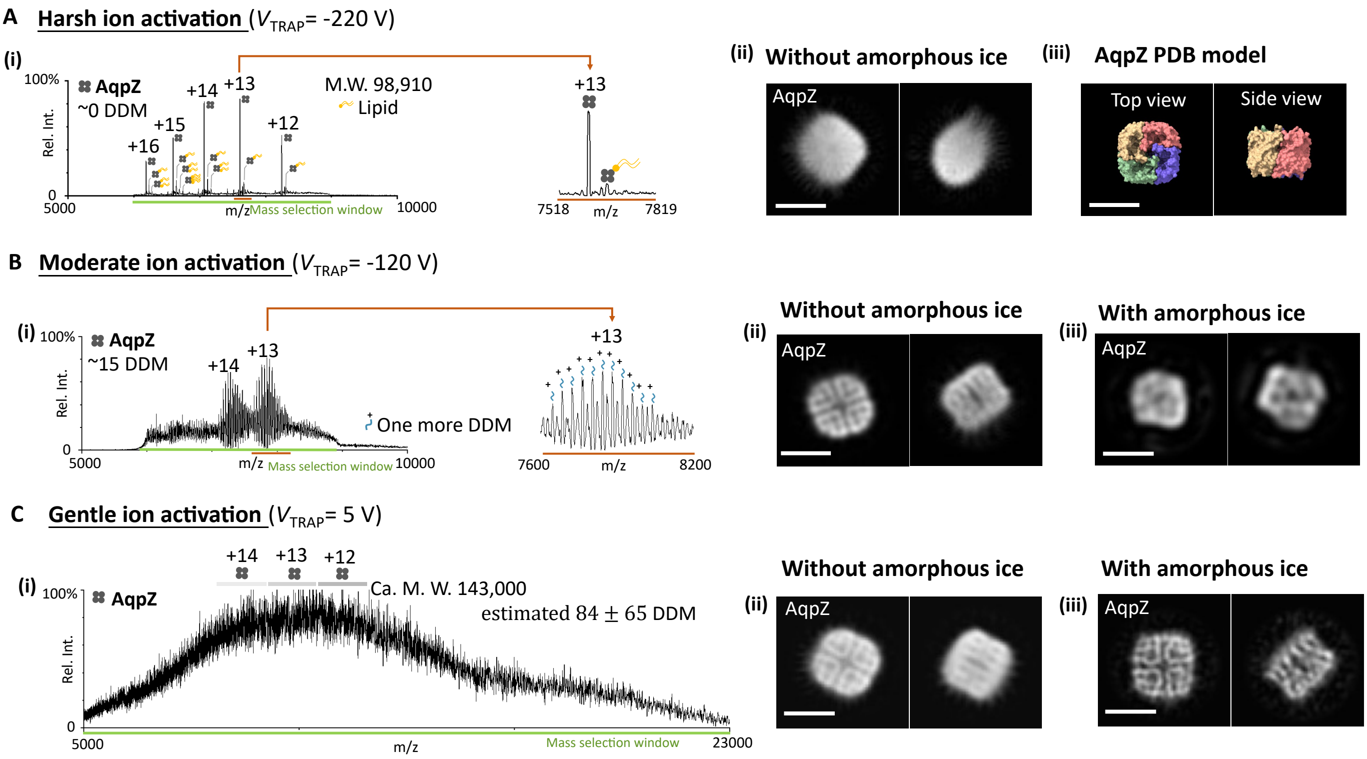


**Fig. S3.**

**Activation-dependent detergent binding in mass spectra of AqpZ, and the corresponding cryoEM 2D classes acquired with and without growth of amorphous ice. (A-C)** Mass spectra and corresponding cryoEM 2D classes of AqpZ under varying activation conditions: harsh, moderate, and gentle. Dimensional -PDB entries 2ABM (AqpZ) is used here. (**i**) Mass spectra, representative 2D classes (**ii**) without and (**iii**) with amorphous ice embedding under the same conditions. Detailed 2D class averages can be found in **Figure S4** for reference. All scale bars, 5 nm. Using the same approach in **Figure S5**, we estimate that under gentle activation conditions 84 ± 65 DDM adducts are presented per AqpZ tetramer.

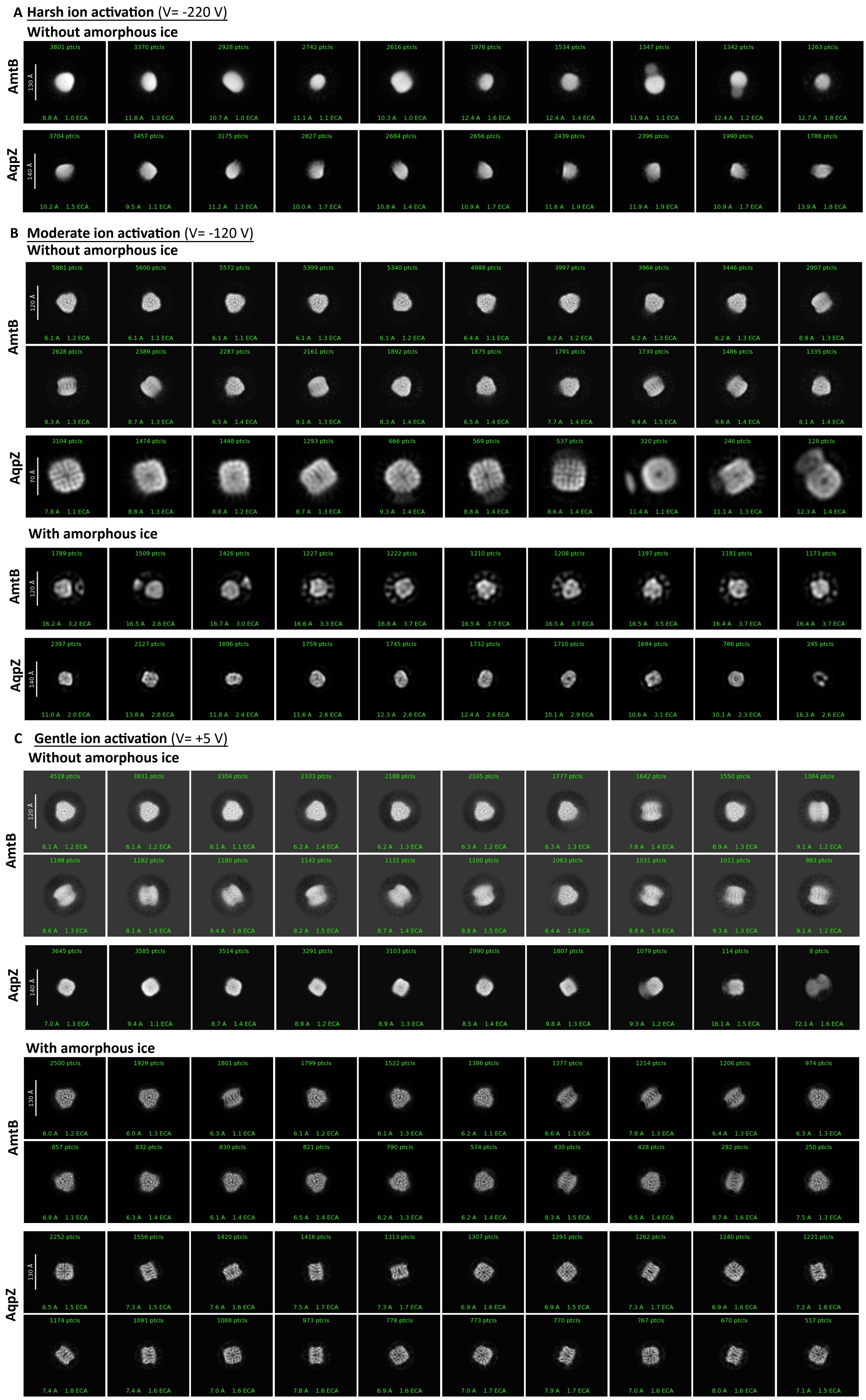


**Fig. S4.**

2D class averages of AmtB and AqpZ under different activation, shown with and without amorphous ice.

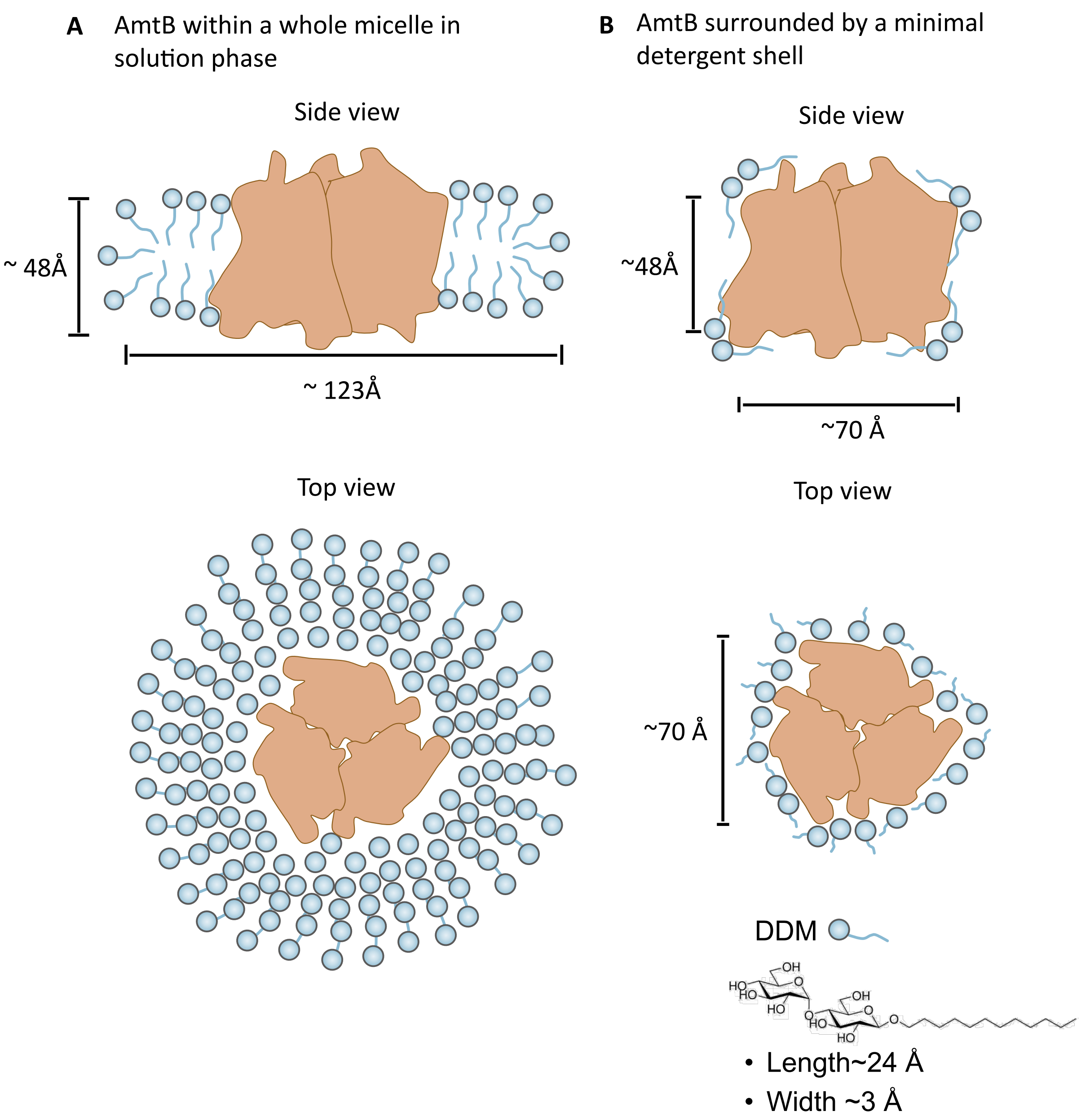


**Fig. S5.**
**Schematic of AmtB detergent encapsulation in solution versus minimal gas-phase conditions (not to scale). (A)** Schematic of AmtB embedded in a full detergent micelle under solution-phase conditions. The micelle forms a continuous shell around the hydrophobic transmembrane region, with an estimated lateral diameter of ~123 Å and a thickness of ~48 Å, based on plunge-frozen cryoEM data. This corresponds to an approximate requirement of ≥500 DDM molecules for complete coverage. **(B)** Schematic of AmtB under gas -phase conditions with partial detergent retention. The micelle is no longer intact, but a minimal shell (no specific interpretation of the DDM molecule) , estimated to involve ≤150 DDM molecules, remains associated with the transmembrane surface. This monolayer-like coverage provides sufficient shielding to preserve structural integrity during gas -phase dehydration and subsequent amorphous ice embedding. Top and side views are shown for both conditions to illustrate the differences in detergent distribution and encapsulation geometry. Schematic illustration of one possible arrangement of the detergent; the exact distribution is not implied.

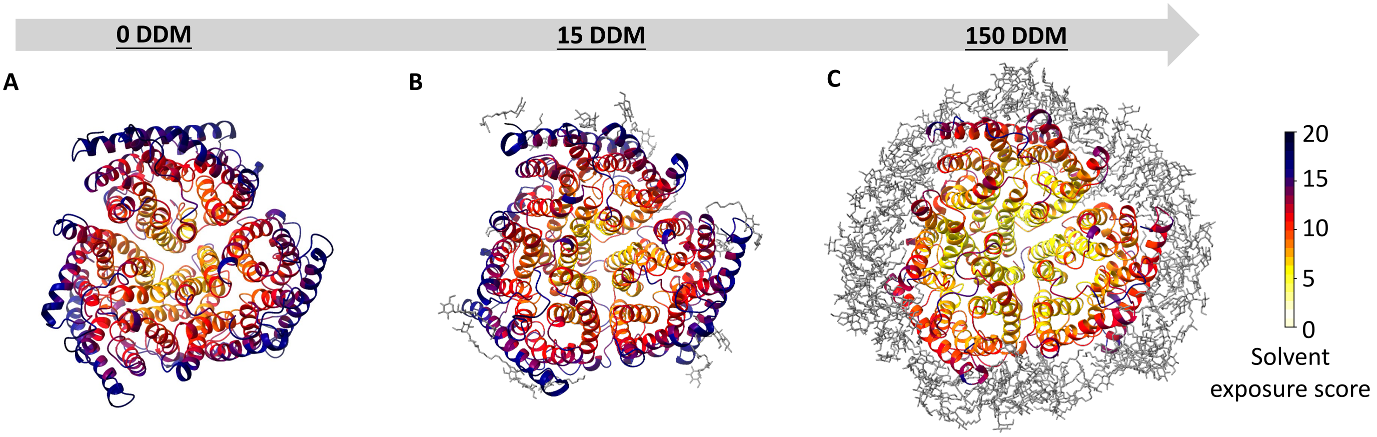


**Fig. S6**

**Solvent-exposure (SE) analysis of AmtB under different detergent conditions.**
SE profiles mirror the per-residue backbone RMSD patterns (light, low SE; dark, high SE). For each residue, SE evaluates the amount and proximity of solvent interactions relative to protein–protein or protein–detergent interactions within a 5-nm cutoff (see SI Methods and Ref (*8*)). High SE values correspond to highly solvent-exposed regions, identifying sites most affected by dehydration. SE mapped onto AmtB models bound to **(A)** 0, **(B)** 15, and **(C)** 150 DDM closely tracks the RMSD-coloured structures. Detergent molecules are coloured grey.

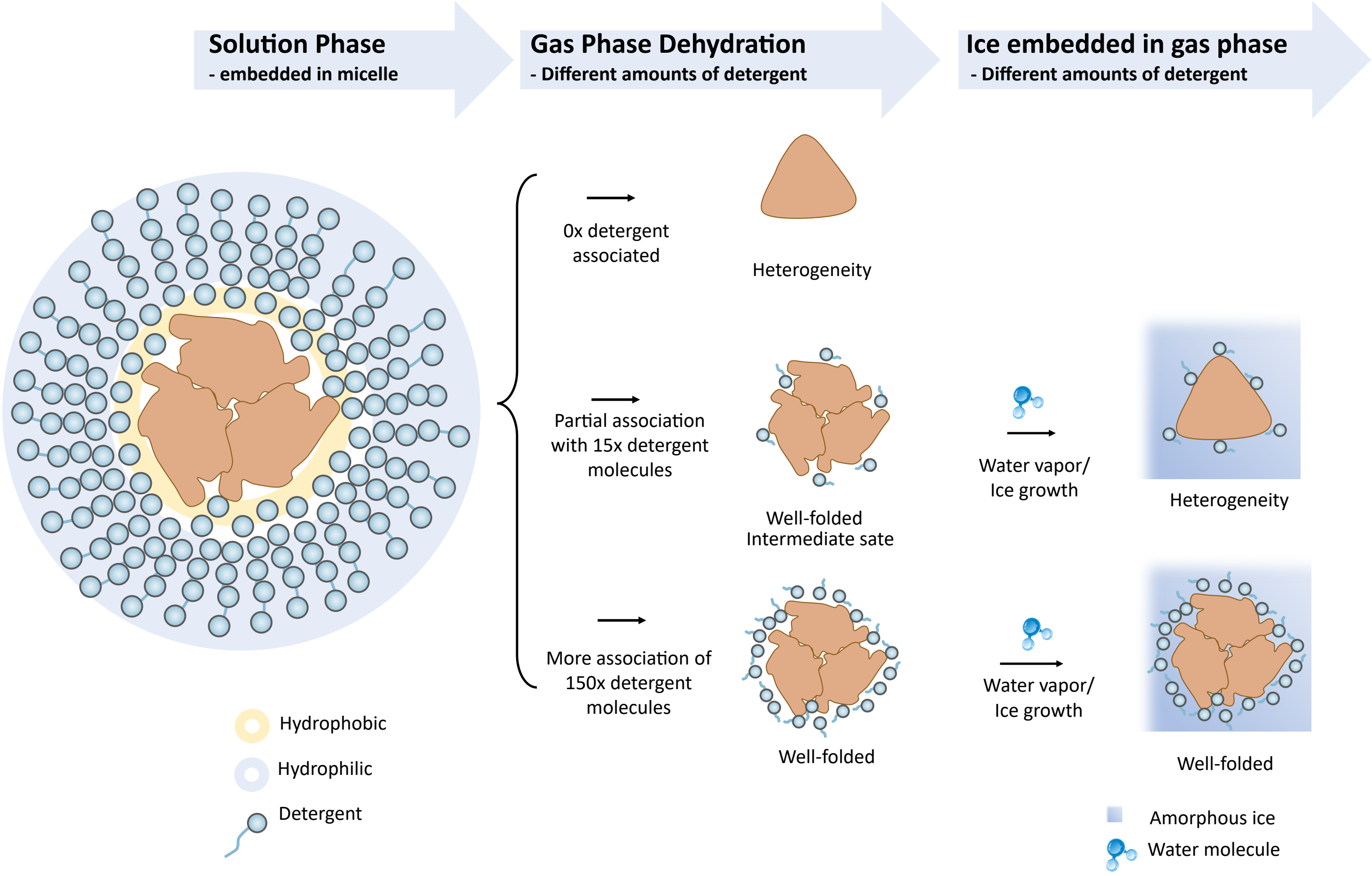


**Fig. S7**

**Proposed mechanism of membrane protein structural preservation during dehydration, detergent removal, and amorphous ice embedding in vacuum.**

Conceptual illustration of the structural response of membrane proteins during the transition from solution to the dehydrated gas phase and subsequent amorphous -ice embedding. Left to right panels depict the solution state (protein in a detergent micelle), varying degrees of detergent retention during gas -phase dehydration, and the rehydrated amorphous -ice state. Three representative detergent association levels are shown: 0 DDM (none), 15 DDM (partial), and 150 DDM (monolayer-like coverage).

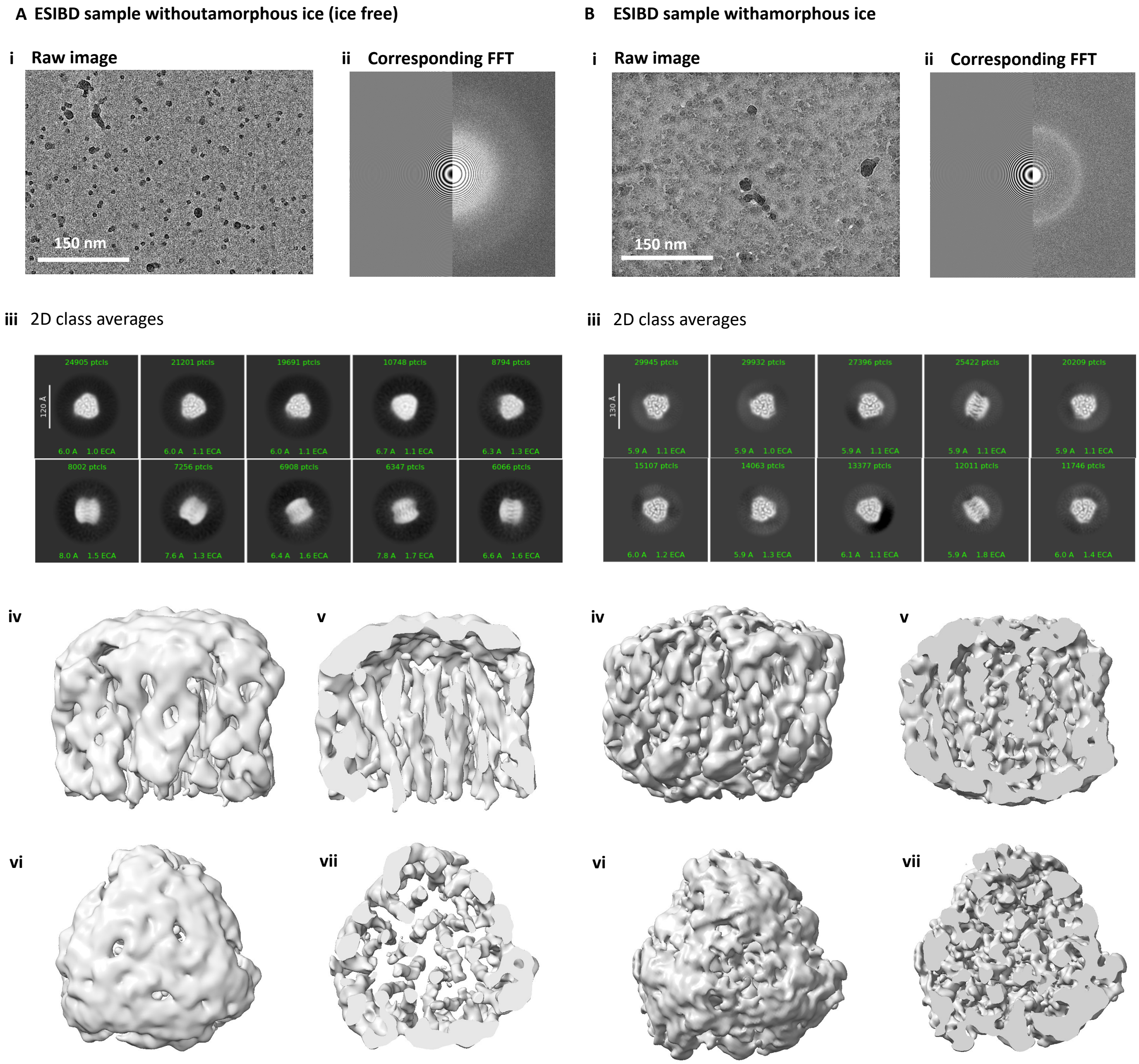


**Fig. S8.**

**CryoEM density map comparison for maps from (A) ice-free and (B) amorphous ice-coated AmtB samples.** The sample without an amorphous ice layer (called ice-free henceforth) showed enhanced particle contrast but failed to yield a high-resolution reconstruction, instead producing a low-resolution envelope with limited internal features and a diffuse peripheral shell. **(i)** Micrographs of AmtB in vacuum after Electrospray ion beam deposition. Without ice coating corresponding power spectrum showing minimal ice, and with ice coating on the protein particle, corresponding power spectrum showing mostly vitreous ice (with occasional crystalline contamination evidenced by scatter diffraction spots in the power spectra). Related FFT figures are shown in **(ii)**. **(iii)** 2D class averages of AmtB. 3D reconstruction map of ice-free AmtB: side view **(iv)** and its cross-section **(v)**, top view **(vi)** and its cross-section **(vii)**. To ensure better comparability between the two results, the ice-free and ice-coated datasets have a similar number of microscope images. The ice-free dataset contains 2382 images, with a total of 198,172 particles used for reconstruction, while the ice-coated dataset contains 2339 images, with 111,218 particles used for reconstruction.

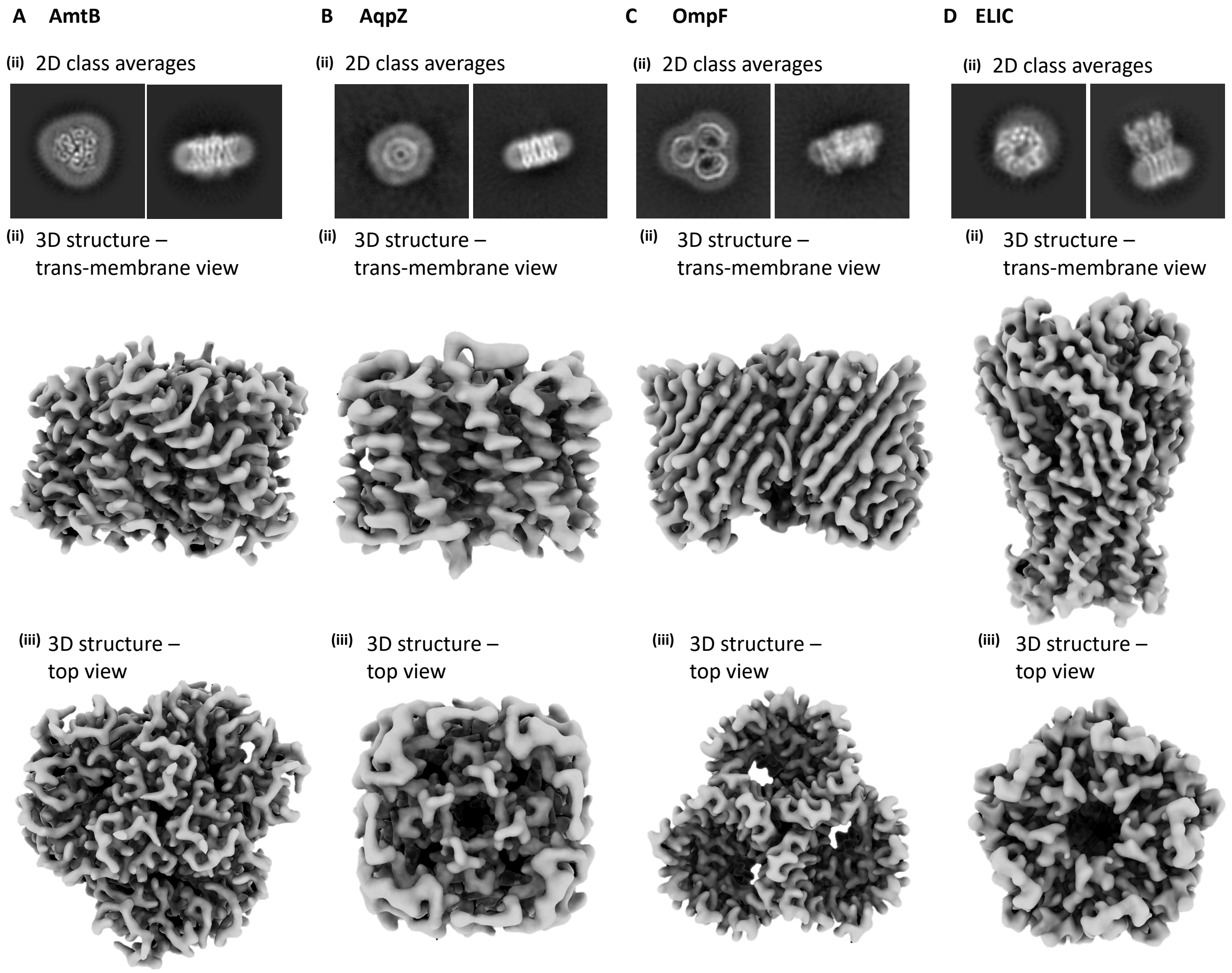


**Fig. S9.**

**Structural integrity and different views of several membrane proteins of plunge -frozen samples (A) AmtB, (B) AqpZ, (C) OmpF and (D) ELIC.** The oligomeric states, representative architecture of all the proteins are well resolved in the plunge-frozen sample: AmtB(trimer, α-helical), AqpZ (tetramer, α-helical), ELIC (pentamer, mixed α-helical and β-sheets with large soluble and TM domains), and OmpF (trimer, β-barrels).

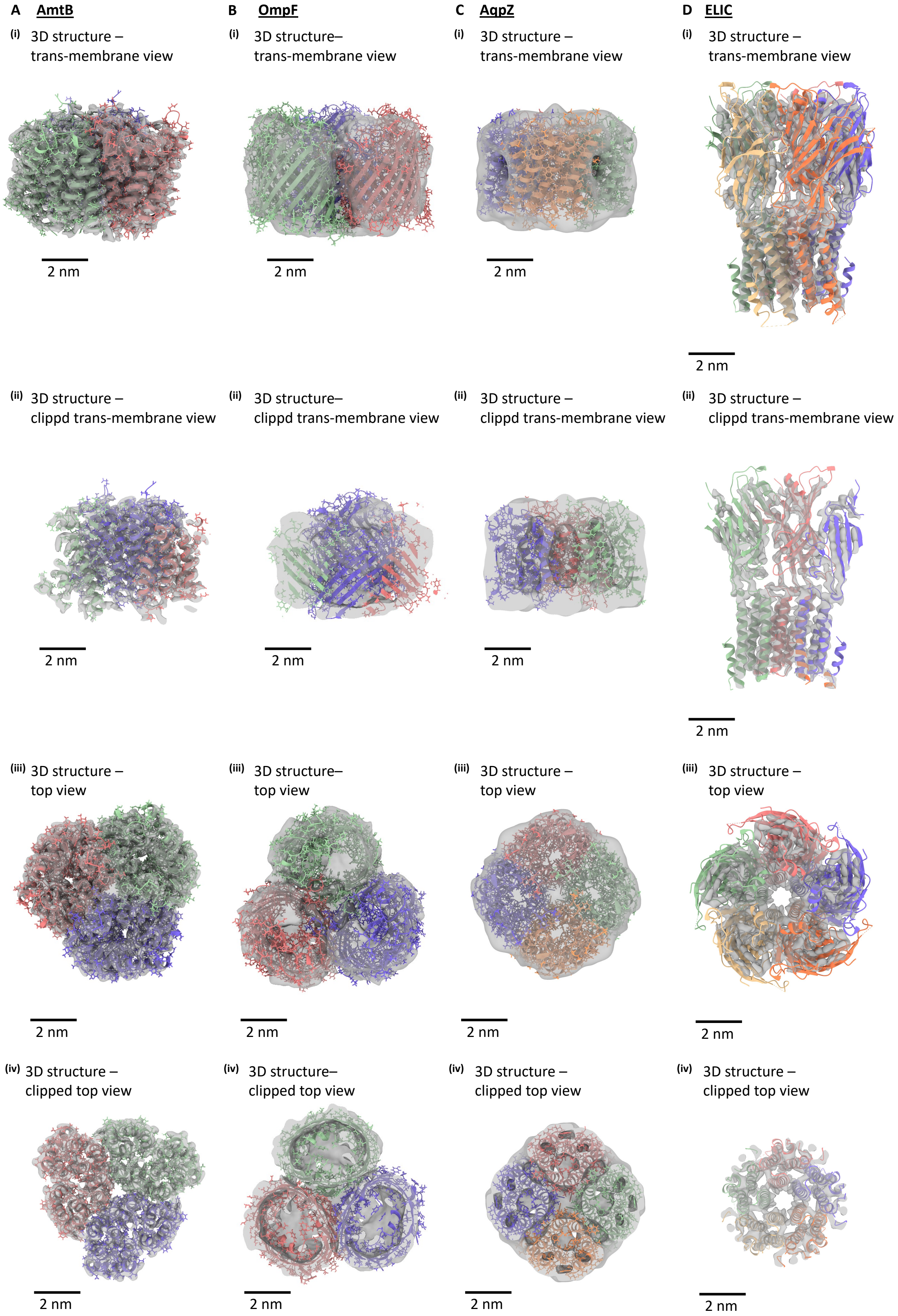


**Fig. S10.**
**Structural integrity and different views of several membrane proteins preserved in vacuum (A) AmtB, (B) AqpZ, (C) OmpF and (D) ELIC.** The maps shown are from vacuum-preserved volumes, whereas the dimensional models were fitted using the corresponding plunge-frozen maps. Scale bars: 2 nm.

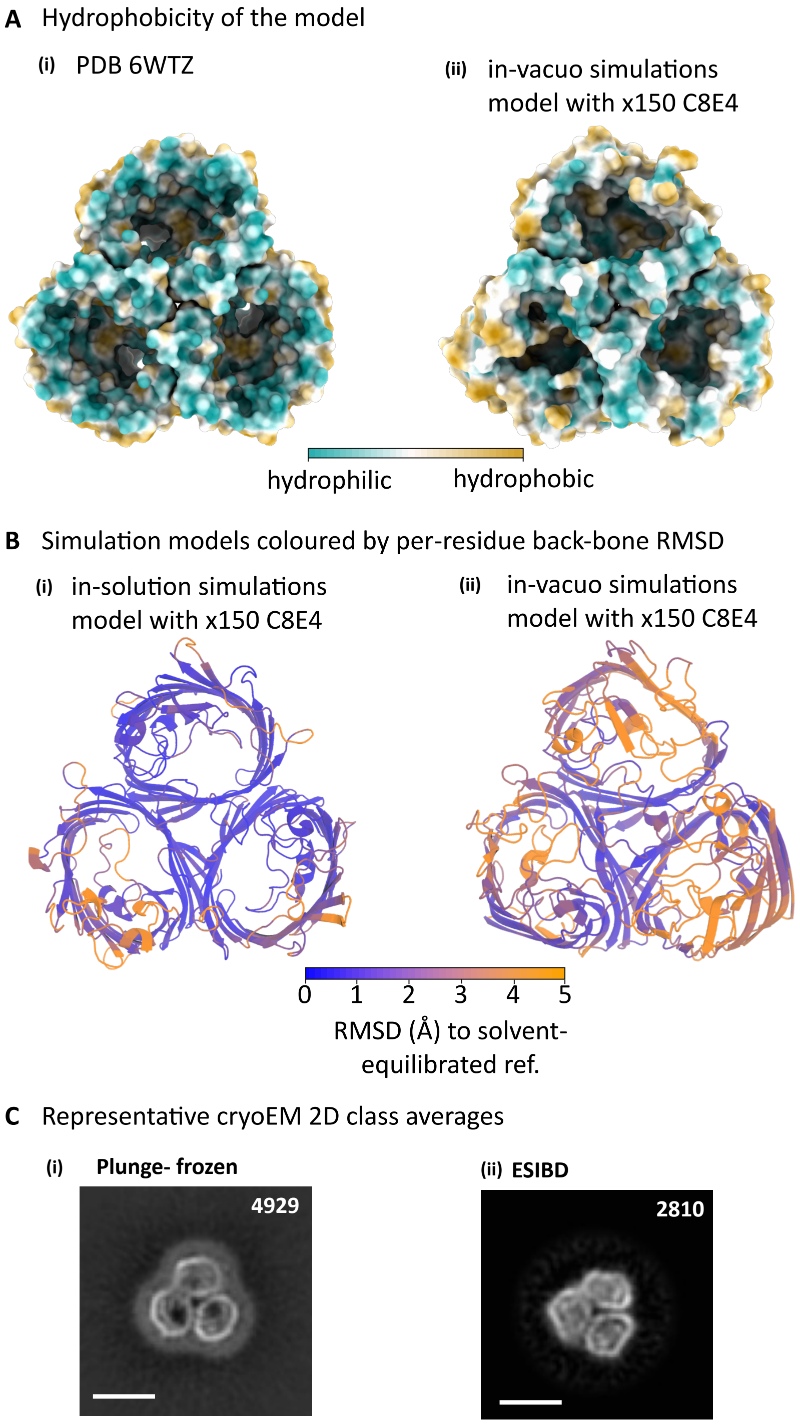


**Fig. S11.**

**MD and cryo -EM evidence for the dented barrel of OmpF. (A)** Hydrophobicity of the models **(i)** Reference (PDB 6WTZ) highlighting the open extracellular and hydrophilic pore. **(ii)** in-vacuo simulations model with 150 C8E4, the barrel dented. (cyan, hydrophilic; yellow, hydrophobic) **(B)** Representative models from the MD simulations coloured by per -residue backbone RMSD (blue, low; orange, high). **(i)** in solution simulation; **(ii)** in-vacuo simulation. The RMSD -coloured model shows elevated deviations consistent with heterogeneity associated with dehydration -induced collapse. **(C)** OmpF 2D class averages. **(i) PF:** pronounced barrel pore shapes. **(ii) ESI–BD:** visibly dented barrel, indicating heterogeneity.

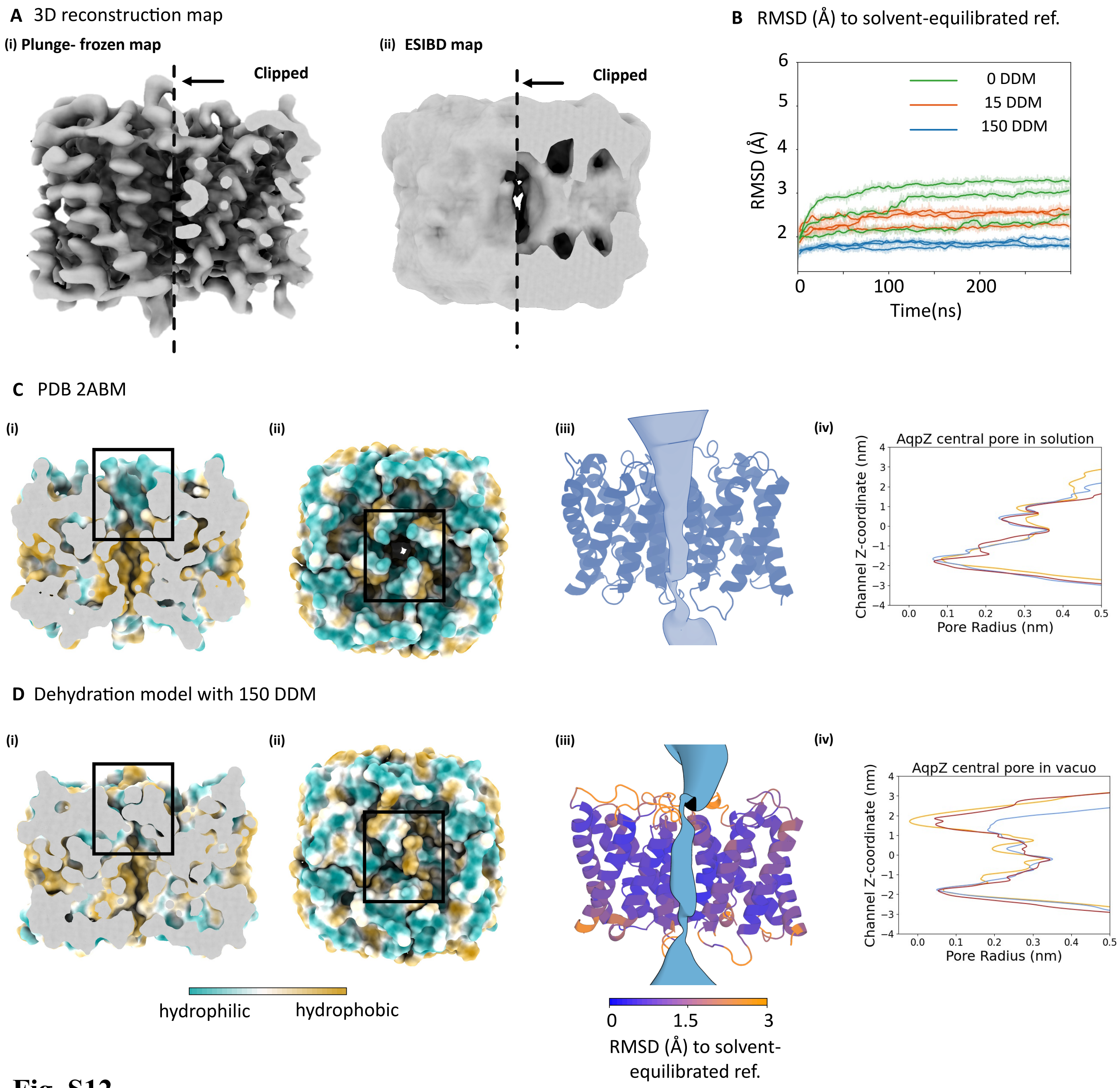


**Fig. S12.**

**MD and cryo-EM evidence for collapse of the AqpZ central pore. (A)** AqpZ reconstructions maps. **(i) PF:** pronounced peripheral helices (unclipped) and ribbon-like pore-proximal helices (clipped). **(ii) ESI–BD:** blurred peripheral helices and re-organised/partially collapsed pore-adjacent helices, indicating heterogeneity in the transmembrane region. (**B**) Backbone RMSD across triplicate trajectories shows minimal RMSD with 150 DDM and larger RMSD with 15 DDM or 0 DDM. **(C)** Central-pore assessments for PDB 2ABM, **(i)**clipped side view (black box) highlighting the open periplasmic pore; **(ii)** top view (black box) of the open pore; **(iii)** CHAP visualization of the pore; **(iv)** pore radius profiles along the z-axis (colours, replicates). **(D)**In the MD dehydration model—even with 150 DDM—the clipped side and top views show periplasmic collapse, which is also evident in the CHAP visualization and pore radius quantification. Together, these indicate a collapsed periplasmic pore in the dehydration model.

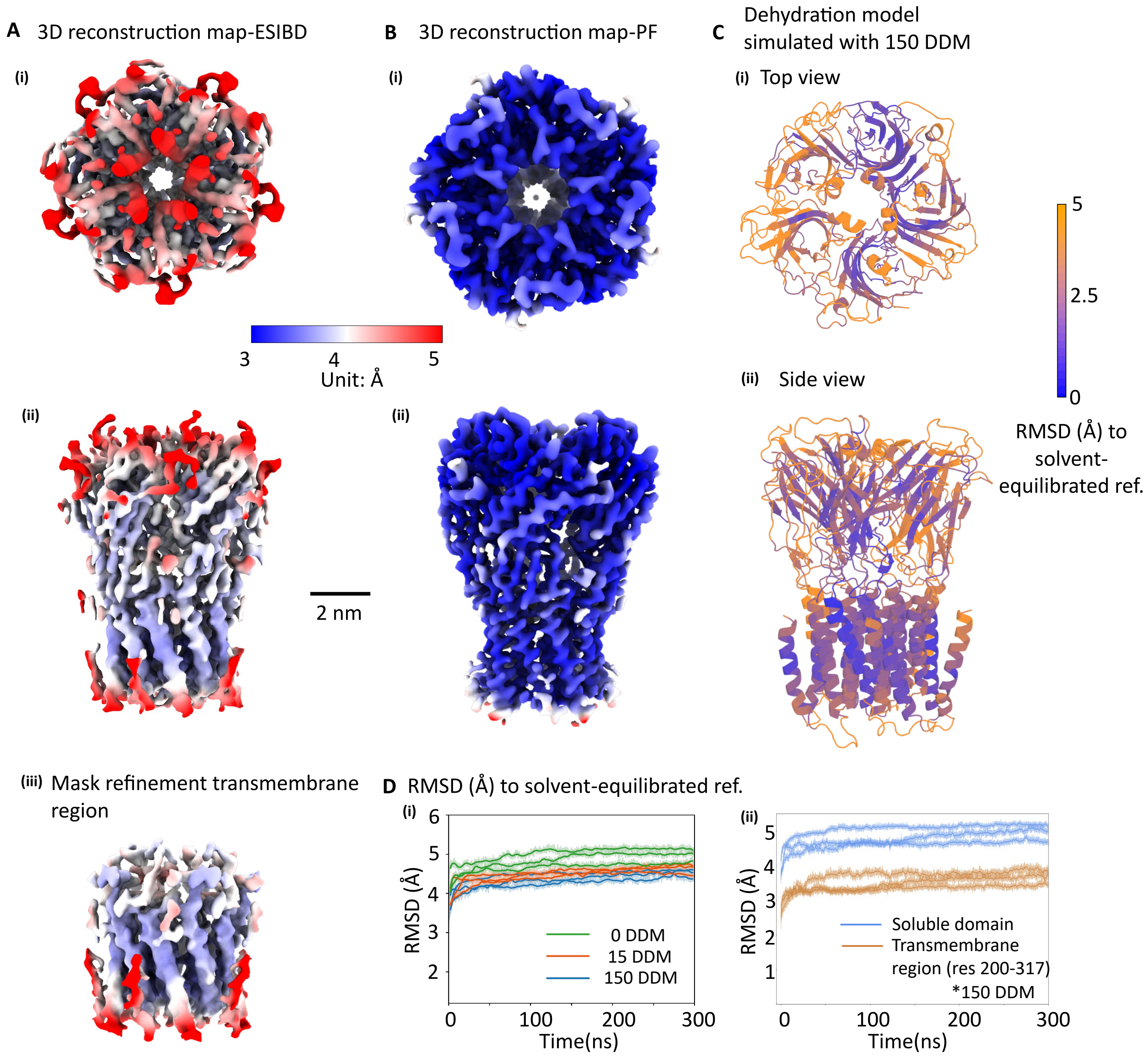


**Fig. S13.**

**MD and cryo -EM evidence for differential preservation of the transmembrane and soluble regions of ELIC.** ELIC cryo-EM maps of **(A)** the ESIBD sample and **(B)** the plunge-frozen sample with local-resolution estimates (blue, high; red, low): **(i)** top view; **(ii)** side view (without mask refinement, Bfactor = 220.1). (A) **(iii)** side view of transmembrane region (with mask refinement, Bfactor = 217.1). The mask was restricted to the transmembrane region, resulting in improved definition of the helices. The TM region of ESIBD map is almost fully resolved (with the exception of a short, flexible helix that is also missing in the PF -control), whereas several peripheral β-sheets and loops are absent in the soluble domain. **(C)** MD dehydration model with 150 DDM retained, coloured by per -residue backbone RMSD (blue, low; orange, high). **(D)** Backbone RMSD across triplicate trajectories: **(i)** whole protein— minimal deviation with 150 DDM and larger deviations with 15 or 0 DDM; **(ii)** separated RMSD traces for the TM and soluble regions highlight differing degrees of preservation. Our MD simulation s match these observations, suggesting that such features are susceptible to dehydration-induced deformation.

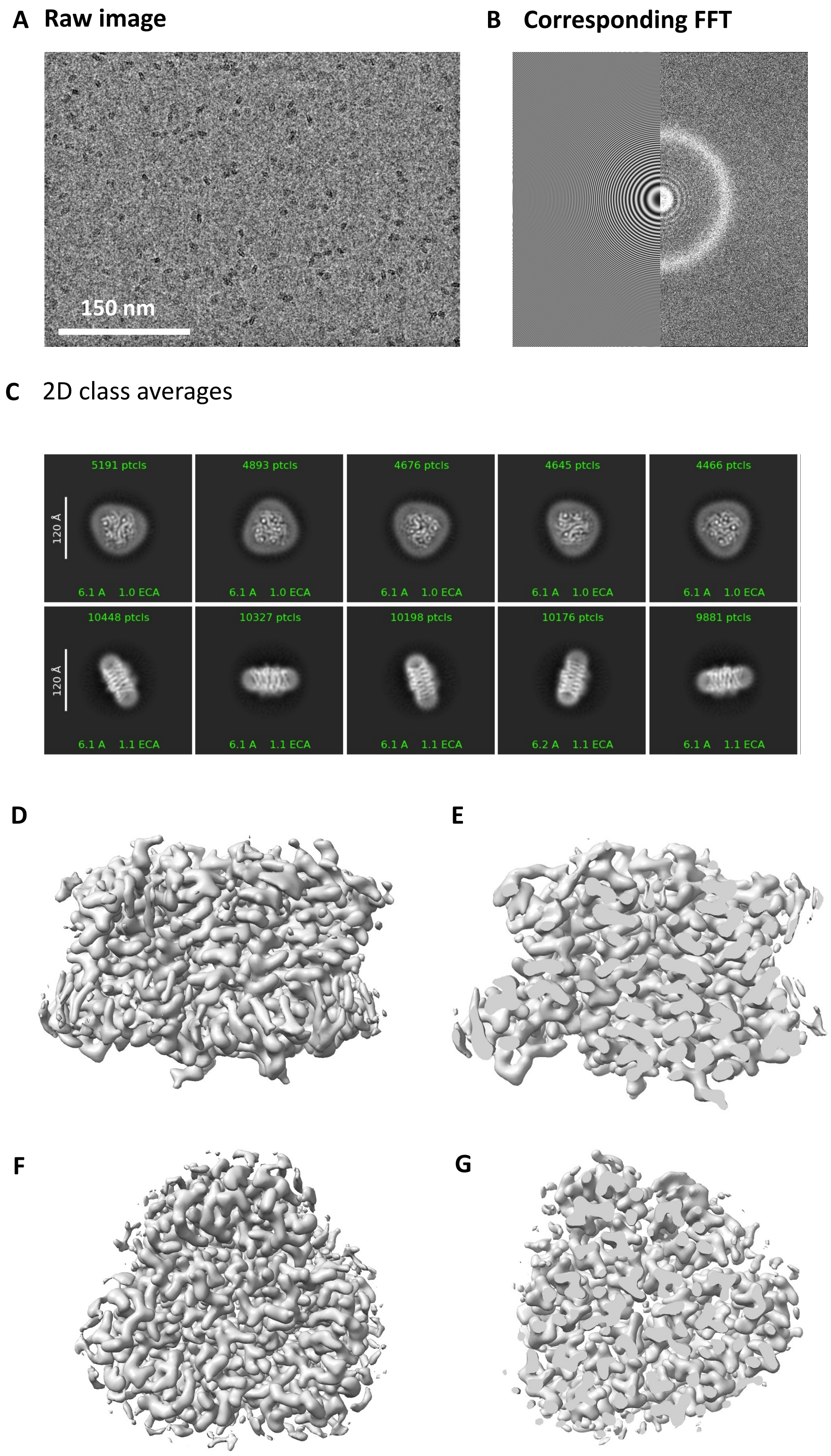


**Fig. S14.**

**Conventional plunge-frozen sample of AmtB with amorphous ice. (i)** Raw micrographs and **(ii)** related FFT. **(iii)** 2D class averages and 3D structure of ice-free AmtB: side view **(iv)** and its cross-section **(v)**, top view **(vi)** and its cross-section **(vii)**.

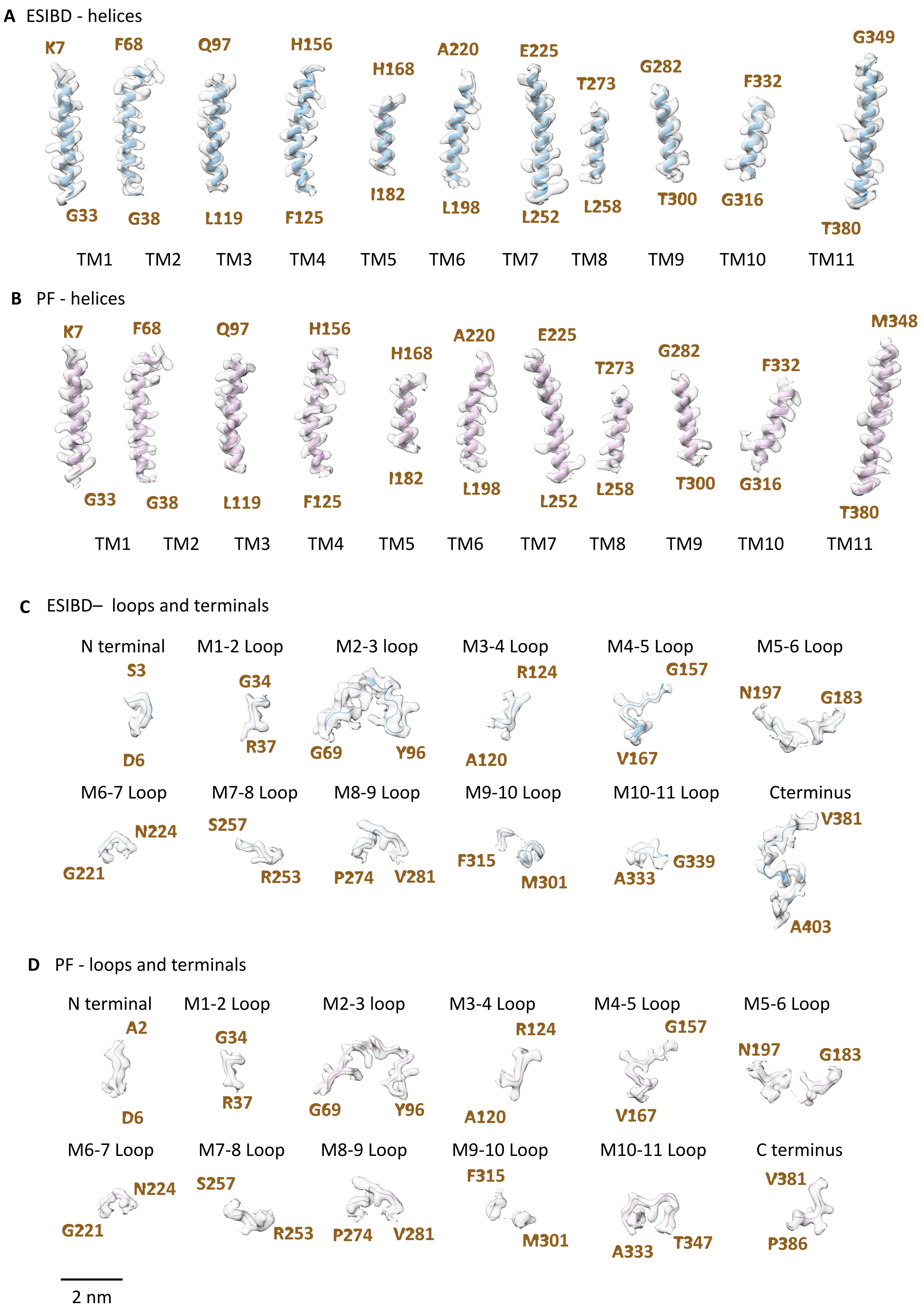


**Fig. S15.**

**Atomic model fitting of AmtB side-chain and backbone density in the cryoEM map via ESIBD method and plunge-frozen method. (A–B)** Representative transmembrane helices (TM1–TM11) shown at B-factor = 0, indicating side chains remain clearly interpretable across most part of the eleven helices, also indicating high-resolution preservation of the transmembrane core. Key residues from each helix are labelled. **(C–D)** Loop and terminal region densities demonstrate clear continuity of the main chain in the loop regions and terminal segments, while some loops exhibit partial disorder without atomic modeling. Loops are labeled according to the helices they connect (e.g., M1–2 loop connects TM1 and TM2).

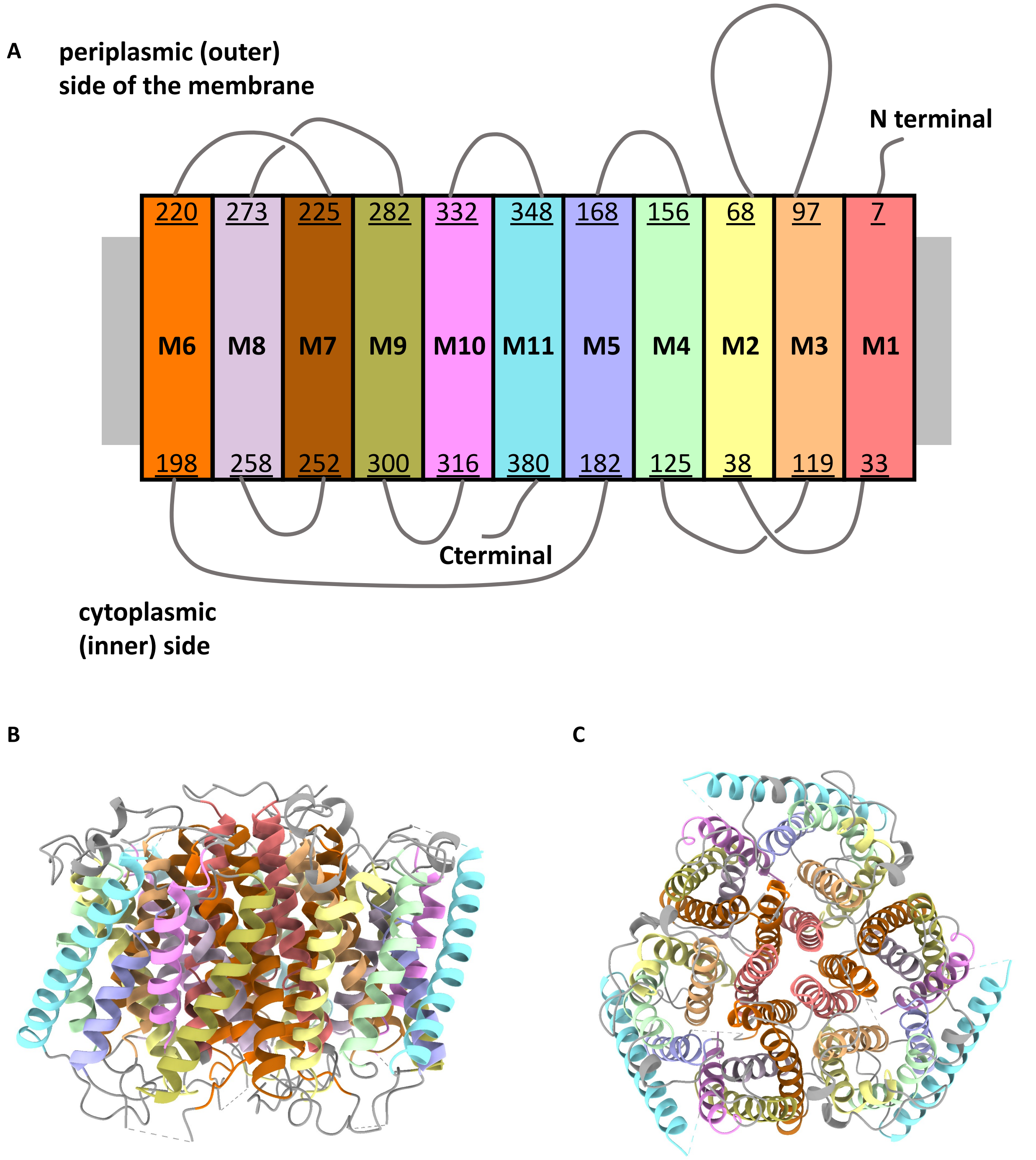


**Fig. S16.**

**Structural organization of the ammonium transporter AmtB.**
**(A)** Topology diagram of AmtB showing the eleven transmembrane helices ( TM1–TM11) colored individually, with N - and C-termini indicated. Periplasmic (outer) and cytoplasmic (inner) sides of the membrane are labeled. **(B)** Side view and **(C)** top view of the trimeric AmtB structure, with each helix colored according to the topology diagram.

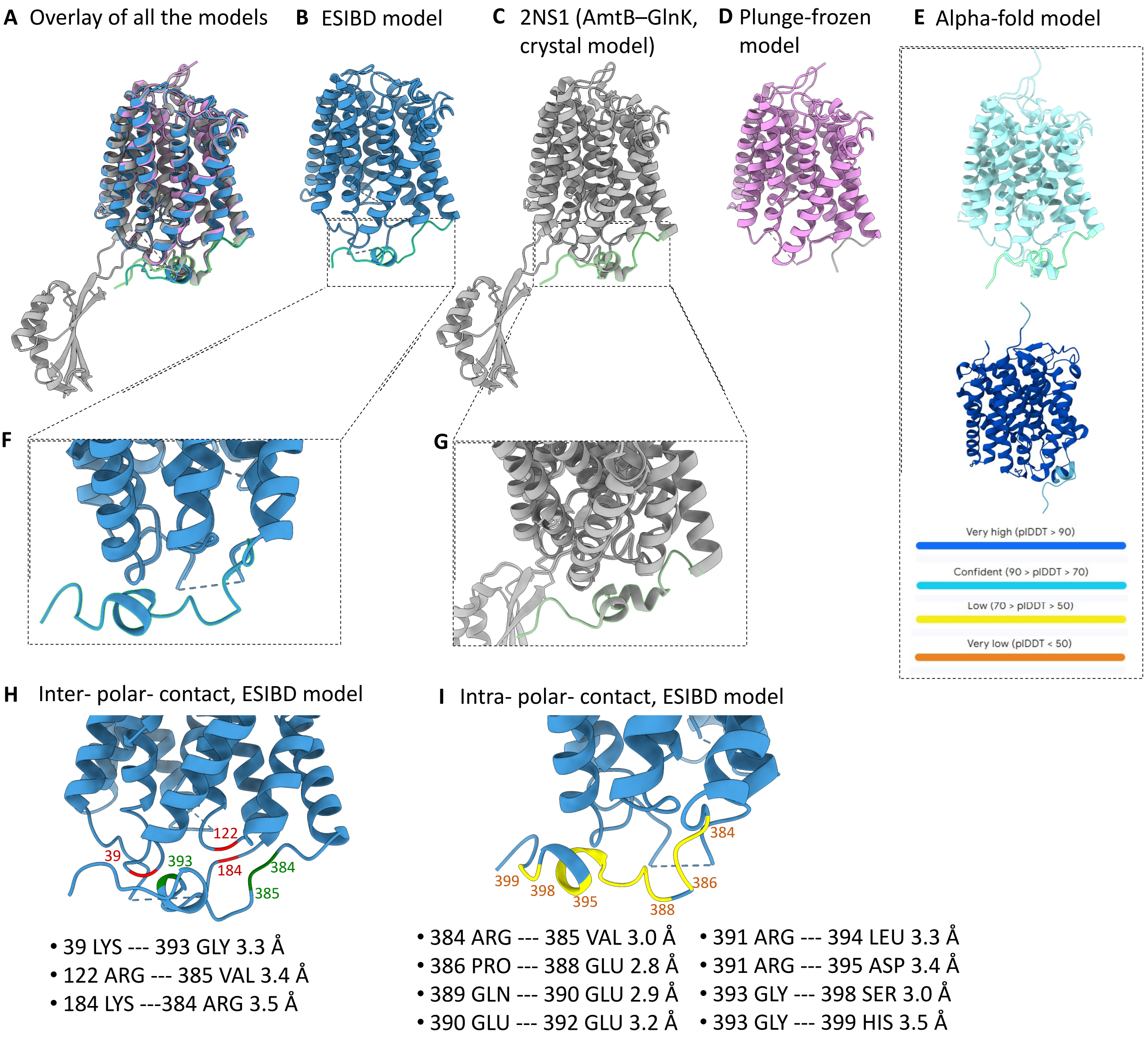


**Fig. S17.**

**Preservation and compaction of the AmtB C terminal under ESIBD.**
**(A)** Overlay of models. **(B)** Cryo-EM model from ESIBD shows a preserved/ordered C-terminal segment (green). **(C)** Crystal structure of E. coli AmtB complexed with the signal-transduction protein GlnK (PDB 2NS1)(*34*). **(D)** In plunge-frozen reconstructions, the distal C terminus is poorly resolved or absent, indicating loss of order upon aqueous vitrification. **(D)** The AmtB model predicted by AlphaFold3 also display similar C terminus.(*35, 36*) **(F)** Close-up of the ESIBD model h ighlights intermolecular/inter-domain polar contacts between the C terminus and nearby cytoplasmic loops that are long -range in solution but become direct in vacuo, consistent with gas -phase compaction, which means this segment can "snap" into a stable helix in a low-dielectric environment. **(G)** Zoomed view of the C terminus in the 2NS1 crystal model. **(H)** Additional ESIBD zoom showing specific inter-loop polar contacts that pin the C terminus. **(I)** Intramolecular contacts within the C terminus, including charge -assisted polar interactions and a protonated histidine ($HisH^{+}$) forming a tight ionic H-bond in vacuum. Together, these observations explain why the C-terminal segment is stabilized in vacuum—by reinforced intra-backbone H-bonds and nearby charge-assisted contacts—yet remains dynamic in solution, consistent with the partly disordered C terminus observe d for the E. coli AmtB – GlnK complex (e.g., PDB 2NS1).

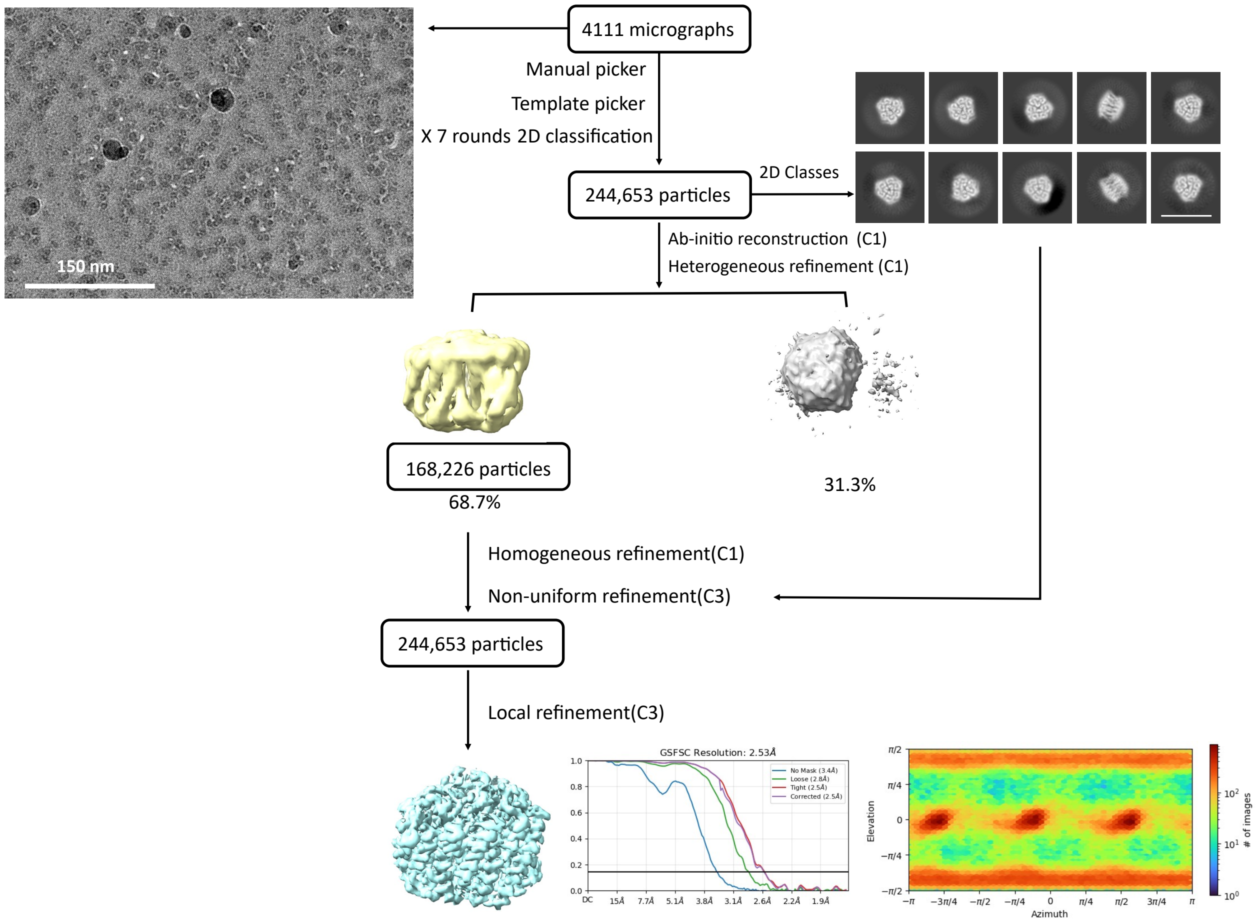


**Fig. S18.**

**Data processing schematic for AmtB with amorphous ice as a native ESIBD sample.**

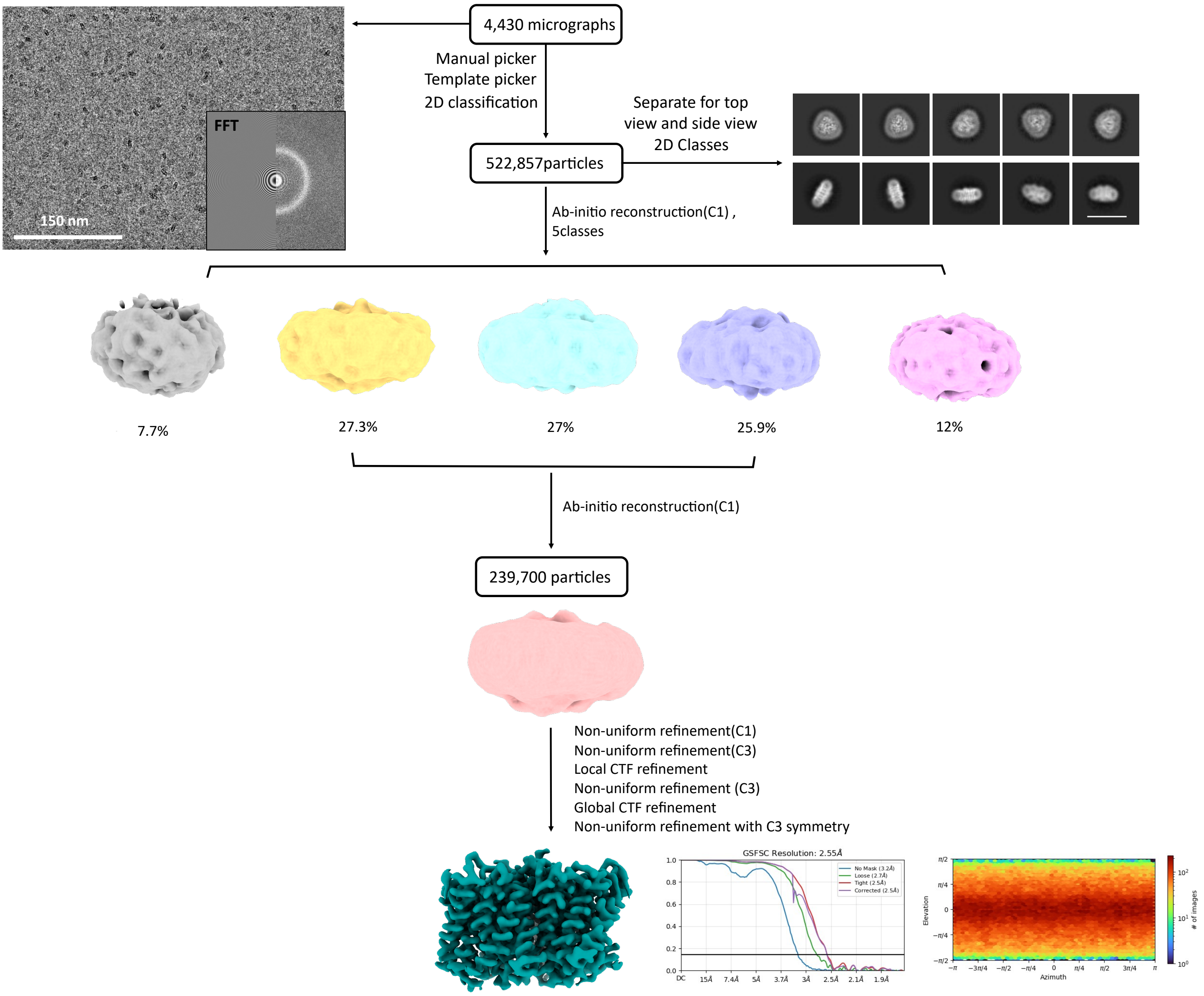


**Fig. S19.**

**Data processing schematic for AmtB from plunge-frozen control sample.**

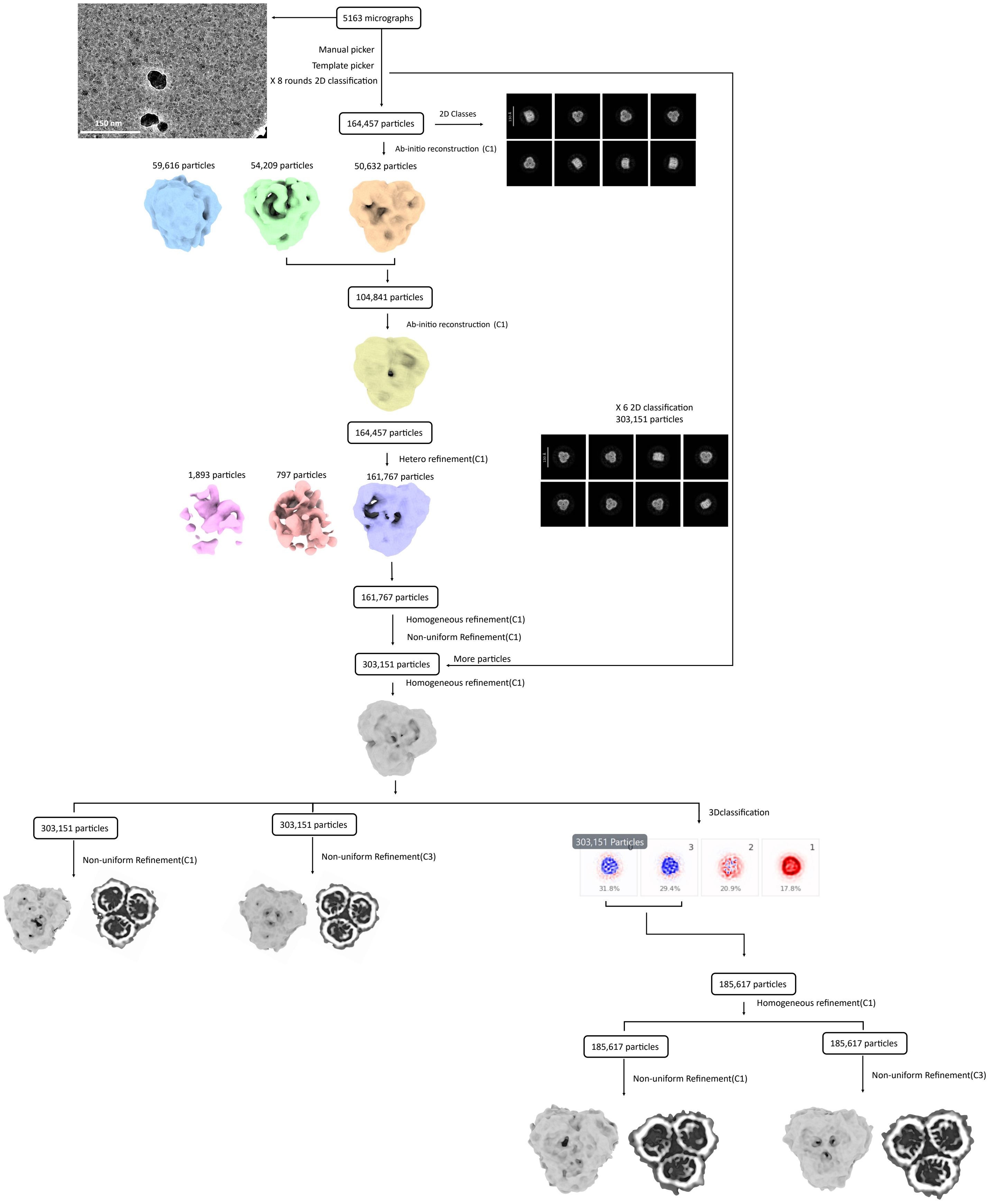


### Fig. S20.

**Data processing schematic for OmpF with amorphous ice as a native ESIBD sample.**

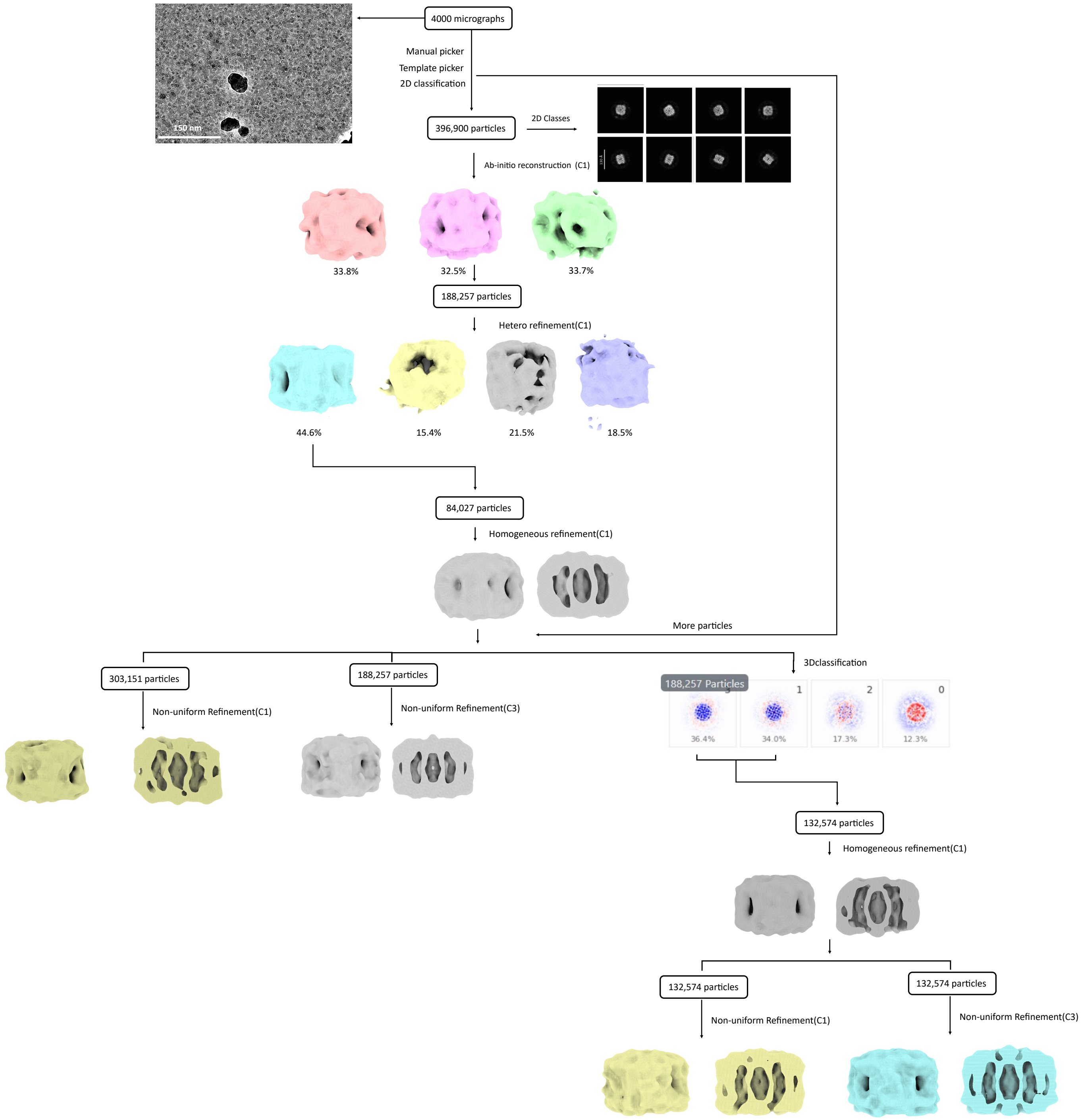


**Fig. S21.**

**Data processing schematic for AqpZ with amorphous ice as a native ESIBD sample.**

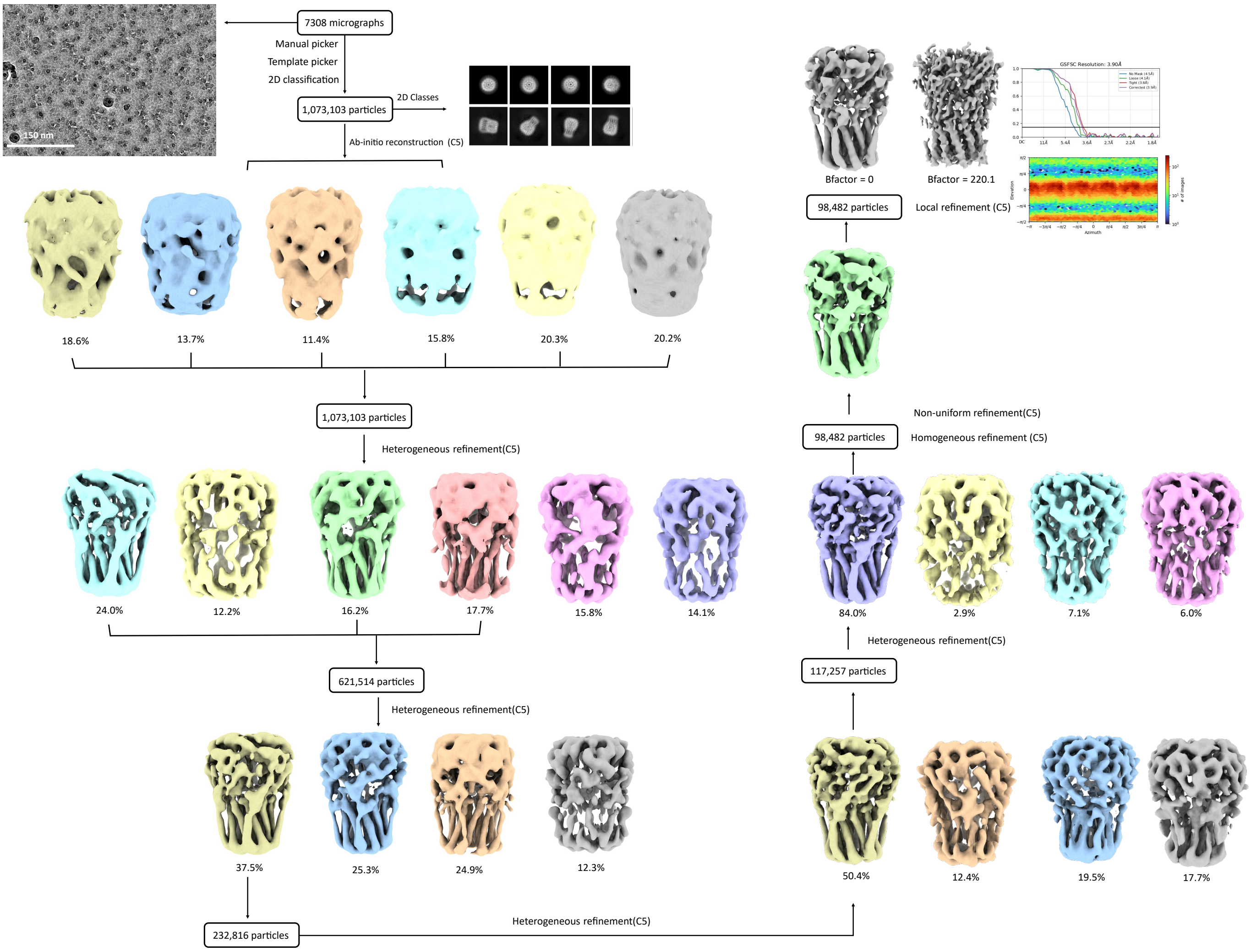


#### Fig. S22.

**Data processing schematic for ELIC with amorphous ice as a native ESIBD sample.**

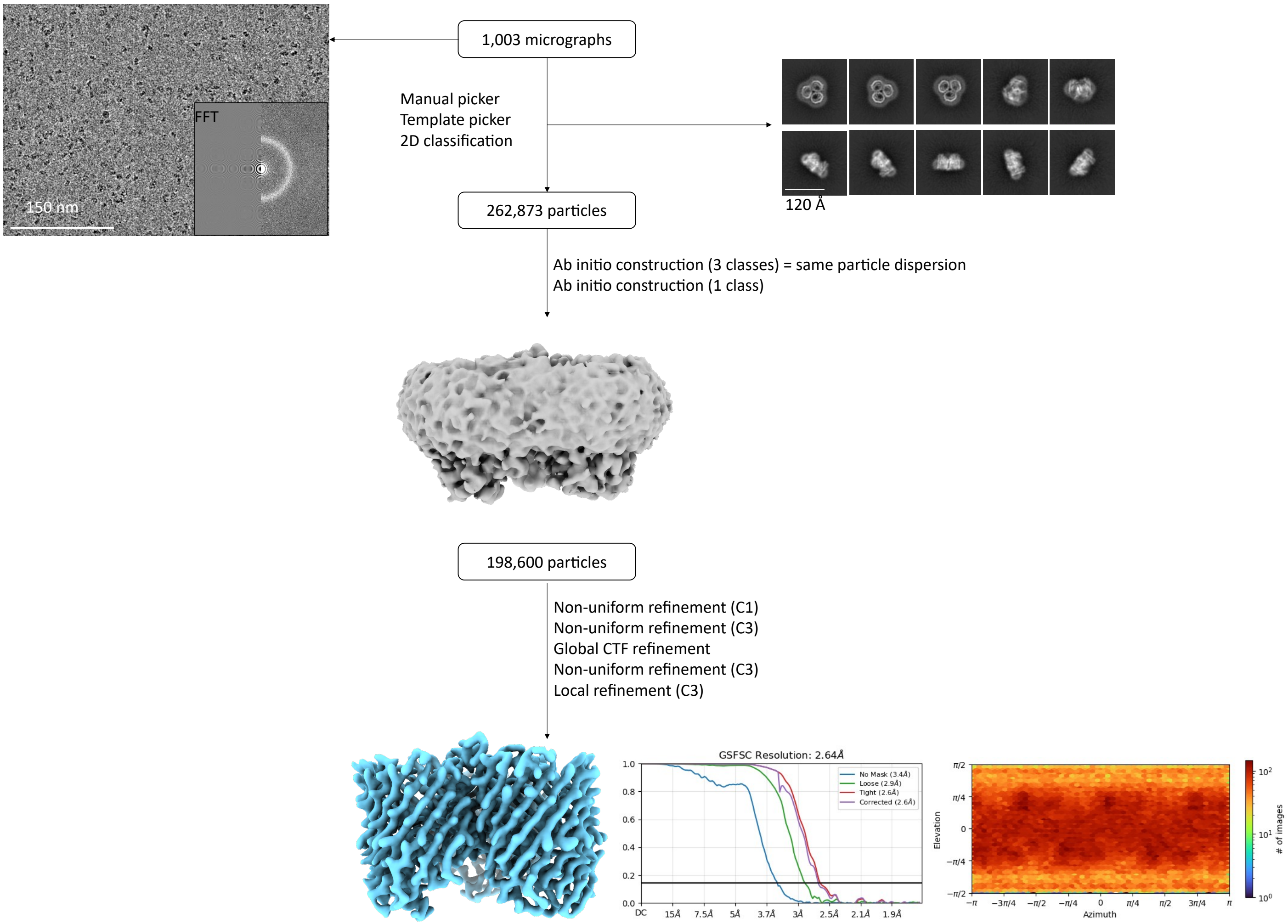


**Fig. S23.**

**Data processing schematic for OmpF with amorphous ice as a plunge-frozen control.**

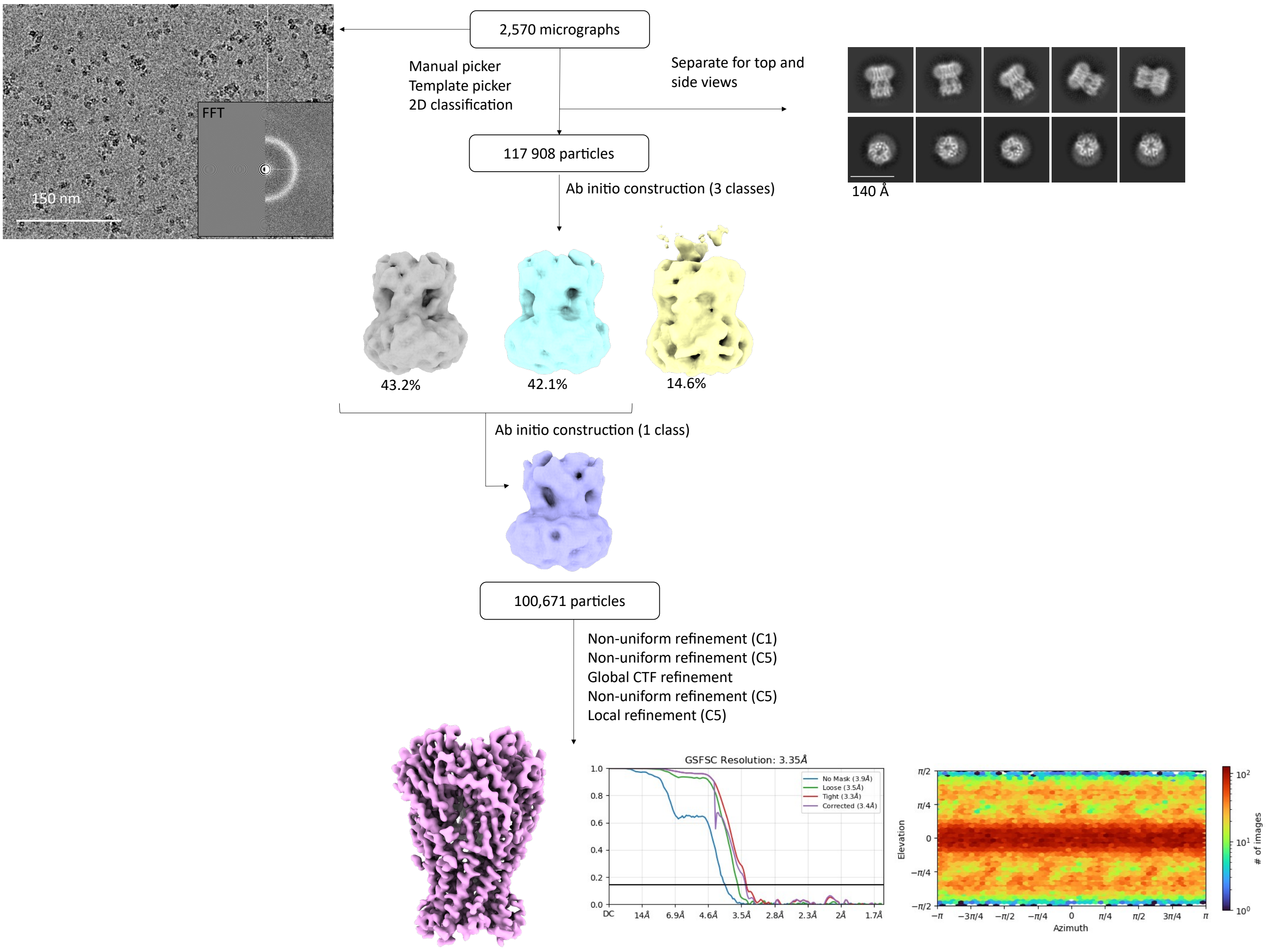


**Fig. S24.**

**Data processing schematic for ELIC with amorphous ice as a plunge-frozen control.**

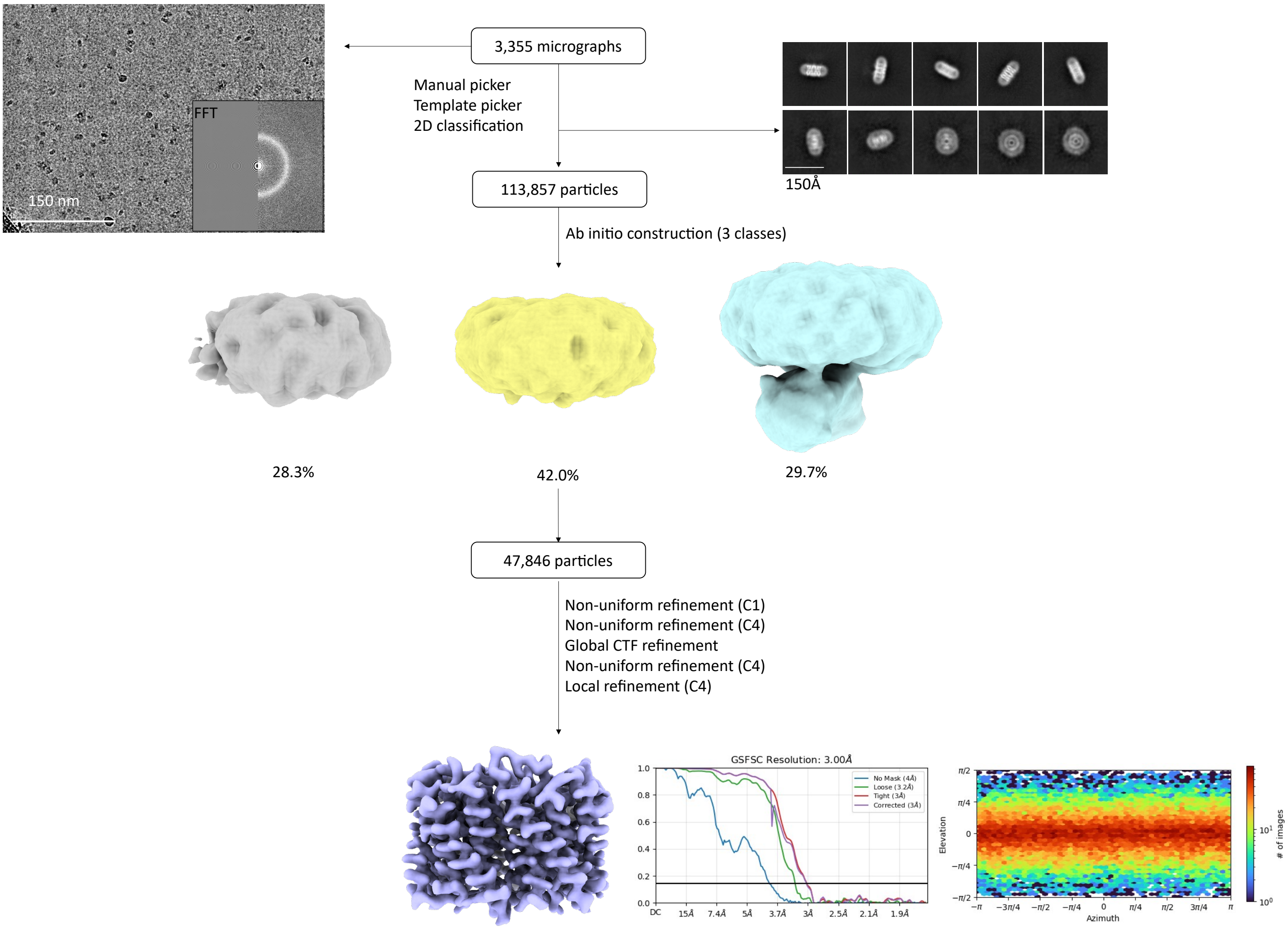


**Fig. S25.**

**Data processing schematic for AqpZ with amorphous ice as a plunge-frozen control.**

**Table S1. The average RMSD values of AmtB model relative to the starting structure for the final 100 ns of MD simulation**

| Condition | Rep1(Å) | Rep2 (Å) | Rep3 (Å) | Mean Value(Å) | SD |
|---|---|---|---|---|---|
| 150 DDM | 3.20 | 3.16 | 3.22 | 3.19 | 0.03 |
| 15 DDM | 4.80 | 4.10 | 4.32 | 4.41 | 0.36 |
| 0 DDM | 5.35 | 4.80 | 5.08 | 5.08 | 0.28 |

**Table S2. Estimated detergent layers for the membrane proteins, calculated using the method shown in Figure S2.**

| | $z_{main}$ | $(\frac{m}{z})_{main}$ | $m_{protein}$ | $m_{detergent}$ | $N_{detergent}$ | $\Delta N_{detergent}$ |
|---|---|---|---|---|---|---|
| AmtB | 16 | 9800 | $126.6\ kDa$ | $510\ Da$ | 60 | 50 |
| ELIC | 30 | 9018 | $185.7\ kDa$ | $510\ Da$ | 166 | 50 |
| AqpZ | 13 | 10900 | $98.9\ kDa$ | $510\ Da$ | 84 | 65 |
| OmpF | 17 | 7700 | $111.3\ kDa$ | $306\ Da$ | 64 | 50 |

**Table S3: Preparation of plunge-frozen control grids.**

| Protein | Working concentration (mg/mL) | Final buffer composition | Grid type | Vitribot Mark IV conditions | |
|---|---|---|---|---|---|
| | | | | Blot force | Blot time (s) |
| AmtB | ~3.2 | 20 mM Tris pH 8, 200 mM NaCl, 0.02% DDM | Quantifoil R1.2/1.3 Cu 300 mesh | 3 | 3 |
| AqpZ | ~1.3-1.5 | 50 mM HEPES pH 7.5, 150 mM NaCl, 0.03% DDM | Quantifoil R2/1 Cu 200 mesh | 3 | 3 |
| ELIC | ~5.2 | 20 mM Tris pH 8, 200 mM NaCl, 0.02% DDM | Quantifoil R2/1 Cu 200 mesh | 3 | 3 |
| OmpF | 1.5 | 20 mM Tris/HCl pH 8.0, 5 mM EDTA, 1% β-OG | Quantifoil R1.2/1.3 Cu 300 mesh | 3 | 3 |

**Table S4. AmtB cryo-EM data acquisition and processing statistics.**

| Data collection and processing | AmtB native ESIBD | AmtB plunge-frozen |
|---|---|---|
| Microscope | Krios G3 | Krios G3 |
| Magnification | 105,000 | 105,000 |
| Voltage(kV) | 300 | 300 |
| Electron exposure ($e^-/Å^2$) | 40 | 40 |
| Defocus range ($\mu m$) | -1.75 to -3 | -1.75 to -3 |
| Pixel size (Å) | 0.85 | 0.83 |
| Symmetry imposed | C3 | C3 |
| Initial particle images (no.) | 435,506 | 522,857 |
| Final particle images (no.) | 244,653 | 239,700 |
| Map resolution (Å) | 2.53 | 2.55 |
| FSC threshold | 0.143 | 0.143 |
| Initial model used (PDB code) | 1U77 | 1U77 |
| Map sharpening B factor ($Å^2$) | 0 | 0 |
| Model composition | | |
| Non-hydrogen atoms | 8454 | 7977 |
| Protein residues | 1152 | 1098 |
| R.m.s deviations | | |
| Bond lengths (Å) | 0.002 (0) | 0.002 (0) |
| Bond angles (°) | 0.467 (0) | 0.374 (0) |
| Validation | | |
| MolProbity Score | 1.44 | 1.17 |
| Clashscore | 3.82 | 3.83 |
| Poor rotamers (%) | 0 | 0 |
| Ramachandran plot | | |
| Favoured (%) | 96.51 | 98.06 |
| Allowed (%) | 3.49 | 1.94 |
| Disallowed (%) | 0 | 0 |

**Table S5. Other MPs via cryo-EM data acquisition and processing statistics.**

| Data collection<br>Processing | OmpF<br>Native ESIBD | AqpZ<br>Native ESIBD | ELIC<br>Native ESIBD |
|---|---|---|---|
| Microscope | Krios G3 | Krios G3 | Krios G3 |
| Magnification | 105,000 | 105,000 | 105,000 |
| Voltage(kV) | 300 | 300 | 300 |
| Electron exposure ($e^{-}/Å^2$) | 40 | 40 | 40 |
| Defocus range ($\mu m$) | -1.75 to -3 | -1.75 to -3 | -1.75 to -3 |
| Pixel size (Å) | 0.85 | 0.85 | 0.85 |
| Symmetry imposed | C3 | C4 | C5 |
| Initial particle images (no.) | 680,960 | 396,900 | 1,073,103 |
| Final particle images (no.) | 303,151 | 188,257 | 232,816 |
| Initial model used (PDB code) | 6WTZ | 2ABM | 6V0B |
| Map sharpening B factor ($Å^2$) | 0 | 0 | 0 |

**Table S6. Other MPs via cryo-EM data acquisition and processing statistics.**

| Data collection Processing | OmpF | AqpZ | ELIC |
|---|---|---|---|
| Microscope | Krios G3 | Krios G3 | Krios G3 |
| Magnification | 105,000 | 105,000 | 105,000 |
| Voltage(kV) | 300 | 300 | 300 |
| Electron exposure ($e^{-}/Å^{2}$) | 40 | 40 | 40 |
| Defocus range ($\mu m$) | -1.75 to -3 | -1.75 to -3 | -1.75 to -3 |
| Pixel size (Å) | 0.83 | 0.83 | 0.83 |
| Symmetry imposed | C3 | C4 | C5 |
| Initial particle images (no.) | 262,873 | 90,890 | 117,908 |
| Final particle images (no.) | 198,600 | 47,846 | 100,671 |
| Map resolution (Å) (masked) | 2.6 | 3.0 | 3.35 |
| FSC threshold | 0.143 | 0.143 | 0.143 |
| Initial model used (PDB code) | 6WTZ | 2ABM | *De novo* ModelAngelo (*37*) |
| Map sharpening B factor ($Å^2$) | 0 | 0 | 0 |
| Model composition | | | |
| Non-hydrogen atoms | 7881 | 6644 | 11780 |
| Protein residues | 1020 | 916 | 1430 |
| R.m.s deviations | | | |
| Bond lengths (Å) | 0.003 (0) | 0.002 (0) | 0.002 (0) |
| Bond angles (°) | 0.484 (0) | 0.499 (0) | 0.386 (0) |
| Validation | | | |
| MolProbity Score | 1.36 | 1.53 | 0.90 |
| Clashscore | 1.97 | 5.00 | 1.58 |
| Poor rotamers (%) | 0 | 0 | 0 |
| Ramachandran plot | | | |
| Favoured (%) | 94.08 | 96.04 | 98.57 |
| Allowed (%) | 5.92 | 3.96 | 1.43 |
| Disallowed (%) | 0 | 0 | 0 |

#### References


1. E. Reading *et al.*, The role of the detergent micelle in preserving the structure of membrane proteins in the gas phase. *Angewandte Chemie International Edition* **54**, 4577–4581 (2015).
2. P. White *et al.*, Exploitation of an iron transporter for bacterial protein antibiotic import. *Proceedings of the National Academy of Sciences* **114**, 12051–12056 (2017).
3. N. G. Housden *et al.*, Intrinsically disordered protein threads through the bacterial outer-membrane porin OmpF. *Science* **340**, 1570–1574 (2013).
4. A. Laganowsky, E. Reading, J. T. Hopper, C. V. Robinson, Mass spectrometry of intact membrane protein complexes. *Nature protocols* **8**, 639–651 (2013).
5. M. N. Webby *et al.*, Lipids mediate supramolecular outer membrane protein assembly in bacteria. *Science Advances* **8**, eadc9566 (2022).
6. H. Hernández, C. V. Robinson, Determining the stoichiometry and interactions of macromolecular assemblies from mass spectrometry. *Nature protocols* **2**, 715–726 (2007).
7. T. K. Esser *et al.*, Cryo-EM of soft-landed β-galactosidase: Gas-phase and native structures are remarkably similar. *Science advances* **10**, eadl4628 (2024).
8. L. Eriksson *et al.*, High-resolution cryoEM structure determination of soluble proteins after soft-landing ESIBD. *arXiv preprint arXiv:2503.22364*, (2025).
9. P. Fremdling *et al.*, A preparative mass spectrometer to deposit intact large native protein complexes. *ACS nano* **16**, 14443–14455 (2022).
10. T. K. Esser *et al.*, Cryo-EM samples of gas-phase purified protein assemblies using native electrospray ion-beam deposition. *Faraday discussions* **240**, 67–80 (2022).
11. M. J. Peet, R. Henderson, C. J. Russo, The energy dependence of contrast and damage in electron cryomicroscopy of biological molecules. *Ultramicroscopy* **203**, 125–131 (2019).
12. K. Amann-Winkel *et al.*, Water's second glass transition. *Proceedings of the National Academy of Sciences* **110**, 17720–17725 (2013).
13. R. Buchner, J. Barthel, J. Stauber, The dielectric relaxation of water between 0 C and 35 C. *Chemical Physics Letters* **306**, 57–63 (1999).
14. T. K. Esser *et al.*, Mass-selective and ice-free electron cryomicroscopy protein sample preparation via native electrospray ion-beam deposition. *PNAS Nexus* **1**, (2022).
15. E. F. Pettersen *et al.*, UCSF ChimeraX: Structure visualization for researchers, educators, and developers. *Protein science* **30**, 70–82 (2021).
16. P. Emsley, B. Lohkamp, W. G. Scott, K. Cowtan, Features and development of Coot. **66**, 486–501 (2010).
17. P. D. Adams *et al.*, PHENIX: a comprehensive Python-based system for macromolecular structure solution. *Biological crystallography* **66**, 213–221 (2010).
18. K. Jamali *et al.*, Automated model building and protein identification in cryo-EM maps. **628**, 450–457 (2024).
19. S. H. Scheres, RELION: implementation of a Bayesian approach to cryo-EM structure determination. **180**, 519–530 (2012).
20. X. Cheng, S. Jo, H. S. Lee, J. B. Klauda, W. Im. (ACS Publications, 2013).
21. J. Lee *et al.*, CHARMM-GUI input generator for NAMD, GROMACS, AMBER, OpenMM, and CHARMM/OpenMM simulations using the CHARMM36 additive force field. *Biophysical journal* **110**, 641a (2016).

22. S. Jo, T. Kim, V. G. Iyer, W. Im, CHARMM‐GUI: a web‐based graphical user interface for CHARMM. *Journal of computational chemistry* **29**, 1859–1865 (2008).
23. M. J. Abraham *et al.*, GROMACS: High performance molecular simulations through multi-level parallelism from laptops to supercomputers. *SoftwareX* **1**, 19–25 (2015).
24. D. Van Der Spoel *et al.*, GROMACS: fast, flexible, and free. *Journal of computational chemistry* **26**, 1701–1718 (2005).
25. G. Bussi, D. Donadio, M. Parrinello, Canonical sampling through velocity rescaling. *The Journal of chemical physics* **126**, (2007).
26. M. Bernetti, G. Bussi, Pressure control using stochastic cell rescaling. *The Journal of Chemical Physics* **153**, (2020).
27. P. Larsson, R. C. Kneiszl, E. G. Marklund, MkVsites: A tool for creating GROMACS virtual sites parameters to increase performance in all‐atom molecular dynamics simulations. *Journal of computational chemistry* **41**, 1564–1569 (2020).
28. L. Konermann, H. Metwally, R. G. McAllister, V. Popa, How to run molecular dynamics simulations on electrospray droplets and gas phase proteins: Basic guidelines and selected applications. *Methods* **144**, 104–112 (2018).
29. H. J. Berendsen, J. v. Postma, W. F. Van Gunsteren, A. DiNola, J. R. Haak, Molecular dynamics with coupling to an external bath. *The Journal of chemical physics* **81**, 3684–3690 (1984).
30. N. Michaud‐Agrawal, E. J. Denning, T. B. Woolf, O. Beckstein, MDAnalysis: a toolkit for the analysis of molecular dynamics simulations. *Journal of computational chemistry* **32**, 2319–2327 (2011).
31. R. J. Gowers *et al.*, "MDAnalysis: a Python package for the rapid analysis of molecular dynamics simulations," (Los Alamos National Laboratory (LANL), Los Alamos, NM (United States), 2019).
32. A. C. E. Dahl, M. Chavent, M. S. Sansom, Bendix: intuitive helix geometry analysis and abstraction. *Bioinformatics* **28**, 2193–2194 (2012).
33. G. Klesse, S. Rao, M. S. Sansom, S. J. Tucker, CHAP: a versatile tool for the structural and functional annotation of ion channel pores. *Journal of molecular biology* **431**, 3353–3365 (2019).
34. F. Gruswitz, J. O'Connell III, R. M. Stroud, Inhibitory complex of the transmembrane ammonia channel, AmtB, and the cytosolic regulatory protein, GlnK, at 1.96 Å. *Proceedings of the National Academy of Sciences* **104**, 42–47 (2007).
35. J. Jumper *et al.*, Highly accurate protein structure prediction with AlphaFold. *nature* **596**, 583–589 (2021).
36. M. Varadi *et al.*, AlphaFold Protein Structure Database: massively expanding the structural coverage of protein-sequence space with high-accuracy models. *Nucleic acids research* **50**, D439–D444 (2022).
37. K. Jamali *et al.*, Automated model building and protein identification in cryo-EM maps. *Nature* **628**, 450–457 (2024).
